# State-dependent evolutionary models reveal modes of solid tumor growth

**DOI:** 10.1101/2022.08.05.502978

**Authors:** Maya A. Lewinsohn, Trevor Bedford, Nicola F. Müller, Alison F. Feder

**Author notes:** To whom correspondence should be addressed ( / /). These authors jointly supervised this work (Nicola F. Müller, Alison F. Feder).

## Abstract

Spatial properties of tumor growth have profound implications for cancer progression, therapeutic resistance and metastasis. Yet, how spatial position governs tumor cell division remains difficult to evaluate in clinical tumors. Here, we demonstrate that elevated cellular growth rates on the tumor periphery leave characteristic patterns in the genomes of cells sampled from different parts of a tumor, which become evident when they are used to construct a tumor phylogenetic tree. Namely, rapidly-dividing peripheral lineages branch more extensively and acquire more mutations than slower-dividing lineages in the tumor center. We develop a Bayesian state-dependent evolutionary phylodynamic model (SDevo) that quantifies these patterns to infer the differential cell division rates between peripheral and central cells jointly from the branching and mutational patterns of single-time point, multi-region sequencing data. We validate this approach on simulated tumors by demonstrating its ability to accurately infer spatially-varying birth rates under a range of growth conditions and sampling strategies. We then show that SDevo outperforms state-of-the-art, non-cancer multi-state phylodynamic methods which ignore differential mutational acquisition. Finally, we apply SDevo to multi-region sequencing data from clinical hepatocellular carcinomas and find evidence that cells on the tumor edge divide 3-6x faster than those in the center. As multi-region and single-cell sequencing increase in resolution and availability, we anticipate that SDevo will be useful in interrogating spatial restrictions on tumor growth and could be extended to model non-spatial factors that influence tumor progression, including hypoxia and immune infiltration.

## Introduction

Tumors develop and progress via an evolutionary and ecological process wherein cellular subpopulations expand and diversify. Over the course of tumor development, tumor cells acquire genetic mutations and new phenotypes that potentially help them compete for resources and adapt for success in their microenvironment. Understanding this process is critical to predicting clinically significant events such as if, how, and when cells metastasize or develop resistance to therapy.

Although tumor cell growth and success is often attributed to genetic and epigenetic aberrations, an additional important determinant of cell growth is physical location within the tumor. Position governs access to oxygen, nutrients, pro-growth signaling from the stroma, pH, cell-cell interactions and degree of immune exposure, all of which can affect cellular proliferation (***Greenspan, 1972***; ***Freyer and Sutherland, 1986a,b***; ***Ward and King, 1997***; ***Petrulio et al., 2006***; ***Marusyk et al., 2014***; ***Lenos et al., 2018***). Taken together, these effects may combine to create an environment in which cells on the boundary of a tumor have higher growth rates compared to those in the center (i.e., “boundary-driven growth”).

Cancer biologists have long been interested in boundary-driven growth because it changes the evolutionary processes and genetic signatures of tumor progression. The evolutionary impact of boundary-driven growth has been explored via evolutionary theory (***Edmonds et al., 2004***; ***Klopfstein et al., 2006***), microbial experiments (***Hallatschek et al., 2007***; ***Korolev et al., 2012***; ***Gralka et al., 2016***), and decades of cancer computational and mathematical models (***Greenspan, 1972***; **Freyer and Sutherland, 1986a,b**; ***Ward and King, 1997***; ***Waclaw et al., 2015***; ***Sottoriva et al., 2015***; ***Sun et al., 2017***; ***Ahmed and Gravel, 2018***; ***Chkhaidze et al., 2019***; ***Noble et al., 2022***). Such investigations have revealed that boundary-driven growth blunts the efficacy of natural selection in selecting for beneficial (i.e, driver) mutations, and purging slower growing (but potentially drug resistant) lineages (***Kayser et al., 2019***). Boundary-driven growth should also enhance the effectiveness of adaptive therapy (***Bacevic et al., 2017***; ***Strobl et al., 2022***) and cell-cell competition in the tumor interior. Further, such growth patterns should distort our expectations for the neutral variant allele frequency spectrum (***Fusco et al., 2016***), which has been used as a null model for identifying signatures of natural selection (***Williams et al., 2016***), and it has been qualitatively suggested in tumor simulation studies that boundary-driven growth could be misinterpreted as selection on tumor trees (***Chkhaidze et al., 2019***). Therefore, establishing and incorporating these null expectations and models for boundary-driven tumor growth is essential in the context of the increasing interest in applying evolutionary theory to clinical disease, for example, in designing adaptive therapy (***You et al., 2017***), identifying driver events (***Turajlic et al., 2018b***; ***Gerstung et al., 2020***), or estimating timings of metastases (***Yachida et al., 2010***; ***Ahmed and Gravel, 2018***).

An extensive history of clinical and experimental observations supports the importance of boundary-driven growth in tumor populations. These observations date back to the pioneering work of Thomlinson & Gray which first identified necrotic structures with surrounding boundaries of growing cells from histological sections (***Thomlinson and Gray, 1955***) and subsequent cell staining approaches found markers of cell division cluster preferentially on the tumor periphery (***Parkins et al., 1991***; ***Connor et al., 1997***). Similar patterns have been noted in cultured tumor spheroids (***Sutherland and Durand, 1984***; ***Mueller-Klieser, 1987***) and organoids (***Florian et al., 2019***; ***Laurent et al., 2013***). Since then, analysis of both clinical samples - via immunohistochemistry (***Hoefflin et al., 2016***; ***Bastola et al., 2020***), spatial transcriptomics (***Berglund et al., 2018***; ***Wu et al., 2021a,b***) and genetic analysis (***Li et al., 2021***; ***Househam et al., 2022***) - and experimental systems such as fluorescently-tracked xenografts (***Lamprecht et al., 2017***; ***Lenos et al., 2018***; ***van der Heijden et al., 2019***; ***Reeves et al., 2018***), have further supported spatial heterogeneity and preferential expansion on the tumor periphery in some tumors.

However, more recent studies have hinted at more complex modes of clinical tumor growth. ***Househam et al. (2022)*** found that many colorectal tumors showed genetic patterns not consistent with boundary-driven growth, and a recent genetic analysis of renal cell carcinomas found the most recent common ancestors of metastatic lineages in the resected tumor interiors as opposed to the tumor boundaries (***Zhao et al., 2021***). Additionally, experimental evidence suggests that although center-bound cells may experience oxygen and nutrient deprivation, hypoxia-related signaling can be linked to stem-cell like tumor phenotypes with heightened survival, increased survival, and chemotherapy resistance (***Lloyd et al., 2016***; ***Chen et al., 2018***). These observations highlight that higher proliferation on the tumor edge is not necessarily synonymous with long-term lineage survival and progression (***Karras et al., 2022***).

A primary challenge in reconciling these conflicting observations is that clinical sequencing often only captures a limited snapshot of tumor diversity and growth. However, this sampled tumor diversity still offers a window into past population dynamics via phylogenetic and phylodynamic tools. *Phylogenetic* approaches, which reconstruct how cells within a tumor are related, have already proven useful in interrogating cancer evolution – for example, in determining the relative ordering of driver mutations (***Gerlinger et al., 2012***; ***Sottoriva et al., 2013***; ***Kim and Simon, 2014***), detecting parallel evolution of gene hits within a tumor (***Turajlic et al., 2018a***; ***Turati et al., 2021***), and resolving whether metastases emerge early or late in tumor development (***Leung et al., 2017***; ***Hu et al., 2019***). In contrast, *phylodynamic* methods, which link shapes of phylogenetic trees to underlying population dynamics, have only rarely been used in cancer genomics (***Alves et al., 2019***), despite widespread application in other fields (***Stadler et al., 2021***; ***Attwood et al., 2022***).

Although phylodynamic approaches have high potential impact in cancer clinical settings, they are generally not adapted to study tumor biology or incorporate the complexities of cancer’s spatial growth. To bridge this gap, we set out to develop a phylodynamic model suited for detecting boundary-driven growth in tumors. First, we quantify characteristic branching and genetic patterns in tumor trees simulated under boundary-driven growth and demonstrate that these patterns correspond to cellular lineages spending different amounts of time on the faster growing tumor edge versus in the tumor center. To fully exploit these patterns for inference, we develop a novel phylodynamic tool based on the multi-type birth-death process (***Maddison et al., 2007***; ***Stadler and Bonhoeffer, 2013***; ***Kühnert et al., 2016***), in which cells have different birth and death rates on the tumor edge and center, and lineages can transition between states as the tumor grows. Crucially, we introduce an extension that links cell birth and mutation, and therefore incorporates rates of sequence evolution that depend on each cell lineage’s inferred history of spatial locations (i.e. spatial states). We provide this state-dependent evolution model (SDevo) as a package in the popular open source Bayesian software BEAST2 (***Bouckaert et al., 2019***). We show that SDevo substantially improves our ability to infer boundary-driven growth dynamics in simulated tumors compared to non-cancer multi-type birth-death models, and validate this approach across a broad array of biological and sampling conditions, including those encompassing selection for driver mutations, 3-dimensional growth, and clinical sampling strategies. Finally, we apply SDevo to spatially-resolved multi-region sequencing data from hepatocellular carcinomas (***Li et al., 2021***) and estimate that cells in the tumor boundary may have birth rates up to 4-6 times faster than those in the interior. More broadly, SDevo is a general tool for quantifying growth processes linked to any discrete state, and future investigations will expand beyond boundary-driven growth.

## Results

### Boundary-driven growth creates distinct phylogenetic tree structures

In order to characterize signatures of boundary-driven growth in tumor trees, we simulate spatially-constrained growth via a cellular agent-based model in a 2D lattice, following a rich literature of studying cancer dynamics via Eden models (***Ermentrout and Edelstein-Keshet, 1993***; ***Anderson and Chaplain, 1998***; ***Waclaw et al., 2015***; ***Chkhaidze et al., 2019***). Simulated tumors grow from single cells over discrete time steps and gain mutations at cell division. Under spatially-constrained *boundary-driven growth*, a cell can only divide if there is an empty lattice spot in its Moore (8-cell) neighborhood, effectively tying its fitness to neighborhood density (Figure S1). Therefore, extant lineages closer to the tumor periphery have progressively higher mean birth rates than those in the center (Figure 1A). For comparison, we simulated a non-spatially constrained *unrestricted growth* model (Figure 1D), in which all cells can divide regardless of density and push their neighbors to create space.

**Figure 1.**
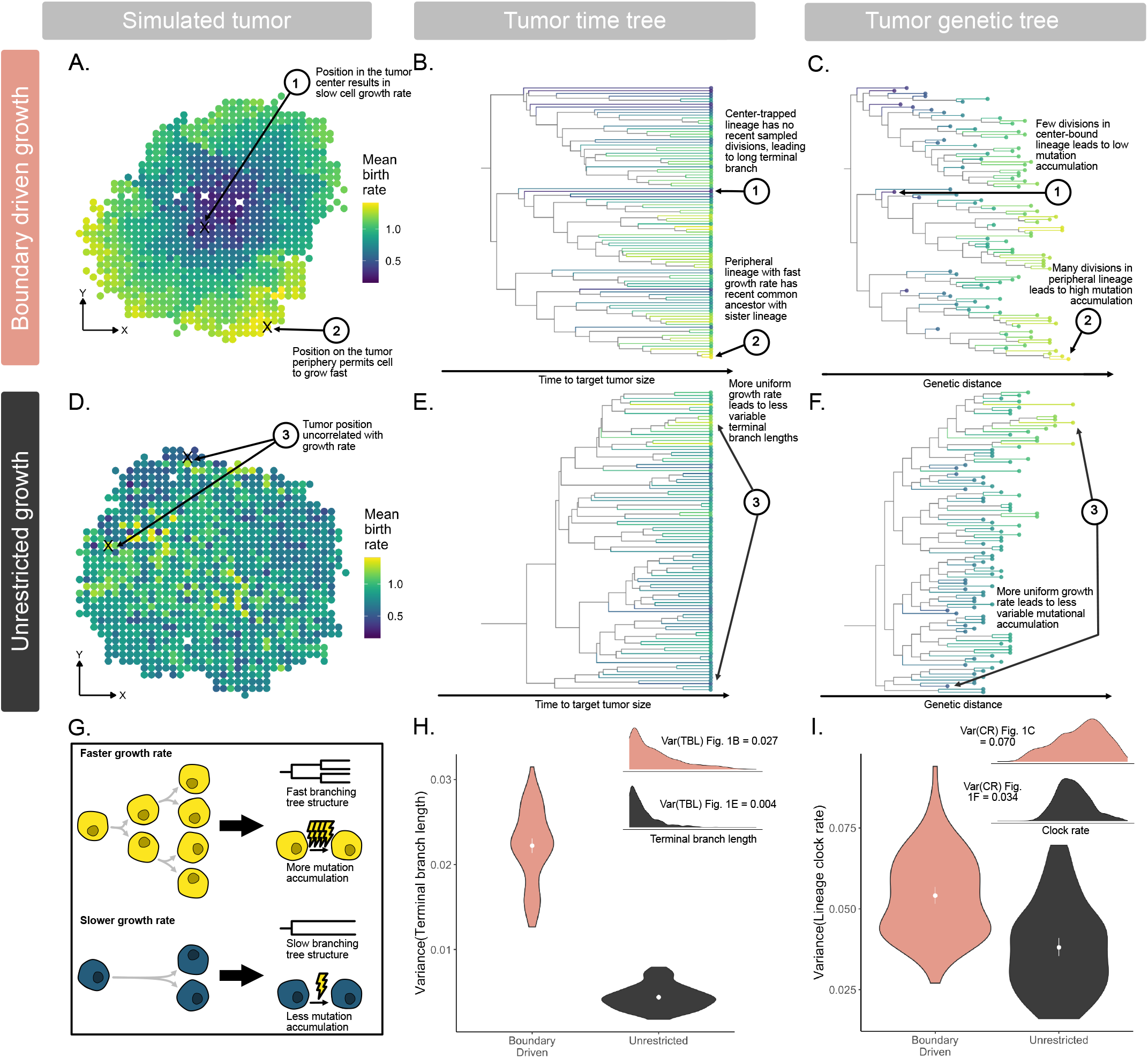
Boundary-driven growth causes characteristic tree patterns associated with asymmetrical division. Representative simulated tumors (α = 0.05) showing variation in mean birth rate in 2D tumor space under **A**. boundary-driven growth via neighborhood-based spatial constraints, and **D**. unrestricted growth. Brighter colors represent higher birth rate (total number of divisions in cell lineage / simulation time) throughout all panels of the figure. **B**. Time tree of the representative boundary-driven growth tumor (subsampled to 100 cells for visualization) shows high variation in branching rates, leading to long terminal branches of center-trapped lineages. **C**. Genetic tree of the representative boundary-driven tumor (subsampled to the same 100 cells) shows ladder-like patterns due to mutation being tied to cell division. **E**. Time and **F**. genetic trees for the representative tumor under unrestricted growth (100 tips visualized) reveal less variation in branching rates and genetic distance. **G**. Cartoon schematic of the two signals of boundary-driven growth in trees left by asymmetric birth rates: variation in branching rates and variation in the number of mutations. Variance in **H**. terminal branch length (TBL) and **I**. clock rate (CR) in tumors under boundary-driven growth and unrestricted growth trees built from all extant tumor cells. Insets show distributions of TBL and CR signals for tumors plotted in **A**. and **B**. Violin plots summarize statistics across 100 simulated tumors.

We first investigated how such growth processes affect the shape and structure of cancer phylogenetic trees to identify detectable tree signals of boundary-driven growth. We consider two types of tree representations – 1) time trees (Figure 1B and E), where the branch lengths are in units of simulation time, and 2) genetic trees (Figure 1C and F), where the branch lengths are in units of number of mutations. We first compare the *time tree* of a tumor simulated under boundary-driven growth (Figure 1B) with one simulated with no spatial restrictions (Figure 1D). In the boundary-driven growth tree, we observe certain leaves (cells) with long terminal branches (i.e., cell 1) and other leaves with much shorter terminal branches (i.e., cell 2). These differential terminal branch lengths directly correspond to both mean lineage birth rate and spatial position within the tumor. Intuitively, lineages trapped in dense center neighborhoods (i.e., cell 1, Figure 1A and B) divide slowly and therefore exhibit longer times since diverging from another sampled cell. Conversely, lineages at the tumor boundary (i.e., cell 2) divide rapidly, and are therefore more likely to be recently related to another sampled cell. We quantify terminal branch lengths in the simulated tumor time trees and find that the asymmetries in birth rates due to spatial constraints result in an overall higher variance in terminal branch lengths under boundary-driven growth (Figure 1H) than under unrestricted growth (Figure 1I).

In Figures 1C and 1F, we reconstruct the *genetic trees* from the same boundary-driven and unrestricted tumor simulations. From this representation of the tumor trees, we observe that if mutation is linked to cellular division, then asymmetries in birth rates across tumor space logically correspond to varying rates of sequence evolution (Figure 1G). This leads to repeated ladder-like patterns of genetic divergence that arise across multiple subclades of the boundary-driven growth tree in which fast-dividing cells on the tumor boundary accumulate more mutations than those in the interior (Figure 1C). These patterns are not observed in the unrestricted growth tree (Figure 1F). We quantify these patterns by measuring variance in mean clock rate (defined by total lineage mutations / simulation time) from extant cells in each simulation and demonstrate that clock rate is more variable across trees derived from boundary-driven growth than in trees simulated under the unrestricted growth model (Figure 1I).

### Boundary-driven growth can be modeled using a two-state birth-death process

As tree structures differ between tumors simulated under boundary-driven and unrestricted spatial constraints, we sought a phylodynamic approach that could differentiate between these two growth modes. One such model is the multi-type birth-death model (***Maddison et al., 2007***; ***Stadler and Bonhoeffer, 2013***; ***Kühnert et al., 2016***), which ties differential rates of birth, death, and sampling of lineages to multiple, discrete states. In our simulation studies, we observe that boundary-driven growth can be effectively simplified into two states. We find that the instantaneous cell birth rate under boundary-driven growth is elevated only in cells immediately adjacent to the tumor edge, but is uniformly low in all cells in the interior (Figure 2A). We can further decompose the tree patterns observed in Figure 1 into edge and center-linked dynamics. As shown in the representative tumor from Figure 1A, all edge-associated cells have short terminal branch lengths. Whereas most of the variation in terminal branch length can be attributed to cells in the center and the mean terminal branch length of cells in the center is more than five times that of cells on the tumor edge (Figure 2B). If we trace the lineages of extant cells back to the root, the fraction of time cell lineages spend on the edge is highly correlated with the variation in mean clock rate observed in Figure 1 (Figure 2C, *R*^2^ = 0.63). In other words, the most mutated cells have spent the majority of their lineage history on the tumor edge. Under unrestricted growth (Figure 2D), we observed no difference between edge and center terminal branch lengths (Figure 2E, ratio of center to edge mean terminal branch lengths = 0.98) and lineage time spent in the edge state is not correlated to clock rate (Figure 2F, *R*^2^ = 0.0016).

**Figure 2.**
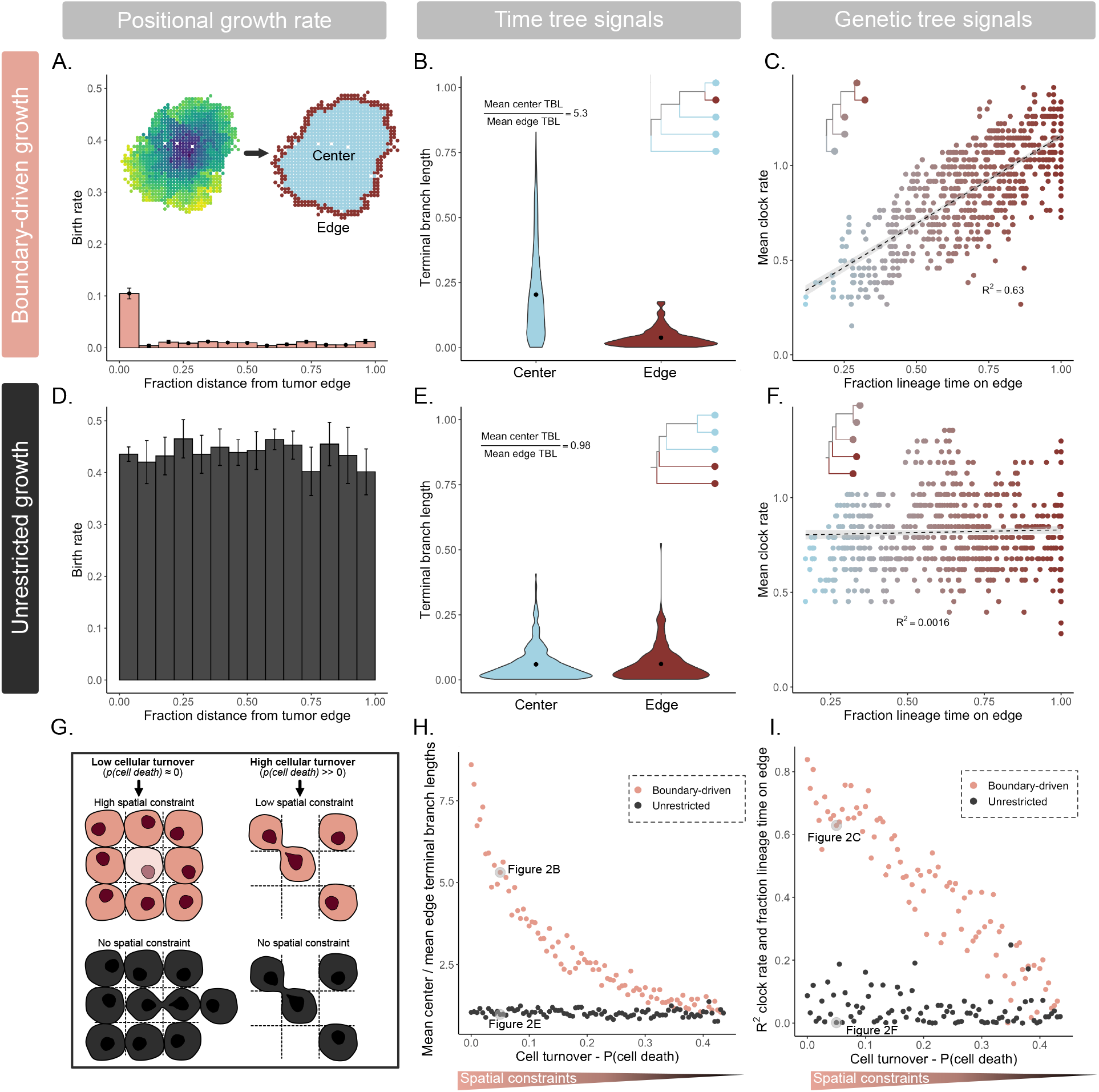
Asymmetries in cell birth rate and signals of boundary-driven growth in trees can be modeled by two-state dynamics. **A**. Histogram of instantaneous cell birth rate as a function of distance from the tumor edge (normalized by maximum distance). Rates are averages over 10 simulations under boundary-driven growth (α = 0.05) with standard error bars. **B**. Distributions of normalized terminal branch lengths (TBL) in a representative tumor under boundary-driven growth (Figure 1A) categorized by leaf edge or center state (inset). **C**. Mean clock rate (total number of mutations / time) of cells in the example boundary-driven tumor versus the fraction of time a cell lineage spends on the tumor edge. Color gradient spans mostly center-associated lineages in blue to mostly edge-associated lineages in maroon. Dashed line is *y* = *x*. **D**. Histogram of instantaneous cell birth rate versus binned distance from tumor edge unrestricted growth simulations (average over 10 simulations with standard error bars, α = 0.05). **E**. Distributions of terminal branch lengths for edge and center leaves in the representative unrestricted tumor (Figure 1D). **F**. Average lineage clock rates versus fraction of time a lineage spends on the tumor edge in the example unrestricted tumor. **G**. Schematic comparing simulated spatial constraints under boundary-driven growth (coral) and unrestricted growth (black). **H**. Ratios of center to edge mean terminal branch lengths across simulations with decreasing spatial constraint (as modulated by cell death rate) under either boundary-driven or unrestricted growth modes. **I**. Correlations (measured via *R*^2^) between fraction of the lineage time spent on the edge and mean clock rate across the same range of spatial constraints and growth modes.

To investigate the robustness of these patterns, we next simulated tumors under a wide range of cell turnover rates. Under boundary-driven growth, increasing cell turnover decreases spatial constraints and therefore lessens the growth advantage between edge and center states (Figure S1, Figure 2G). We measured the ratio of mean center to edge terminal branch lengths as in Figure 2B and E across these different effect sizes and found that this ratio is a consistent indicator of boundary-driven growth that decreases as spatial constraints are relaxed (Figure 2H). The correlation between fraction of lineage time spent on the edge and mean clock rate is also specific to the boundary-driven growth model and sensitive to effect size (Figure 2I). Therefore, we conclude that the patterns left by boundary-driven growth can be effectively approximated by a two-state birth-death model.

### Phylodynamic models can recover signals of boundary-driven growth in tumor phylogenies

Two-state birth-death models incorporate how lineages divide, die, change states, and are sampled. In this class of models, birth events correspond to observed branching events on the tree and the rate of these branching events depends on an underlying type or state. Although existing phylodynamic models, such as BDMM’(***Kühnert et al., 2016***; ***Vaughan, 2022***) and BiSSE (***Maddison et al., 2007***), permit asymmetrical division rates based on state, they do not link birth and mutation. Therefore, although they are well-positioned to infer faster birth rates from branching structure, they cannot learn from differential rates of genetic divergence, a key hallmark of boundary-driven growth we observed in simulations. Additionally, branching patterns are prone to artificial inflation if more cells from a particular state are sampled in a clustered manner (***Höhna et al., 2011***). Thus, existing models both do not incorporate all potential signals (i.e., clock rate differences) and, importantly, may be biased by sampling procedures in clinical tumor biopsies. To address these shortcomings, we introduce a **S**tate-**D**ependent sequence **evo**lution (SDevo) model to directly tie state-dependent birth rate to clock rate. This enables the model to learn from both clock rate and branching rate signals that arise from boundary-driven growth (Figure 3A). SDevo accepts genetic sequences sampled from distinct spatial locations, along with a cell-state label (i.e., center versus edge). It generates posterior distributions of phylogenetic trees alongside joint estimates of phylodynamic model parameters. Inferred trees are time trees, which encompass the order and timing of cellular divergence events and include inferred internal node states, representing the location of unsampled ancestral cells. Model parameters include state-dependent birth and death rates, and the rate at which cells transition between states.

**Figure 3.**
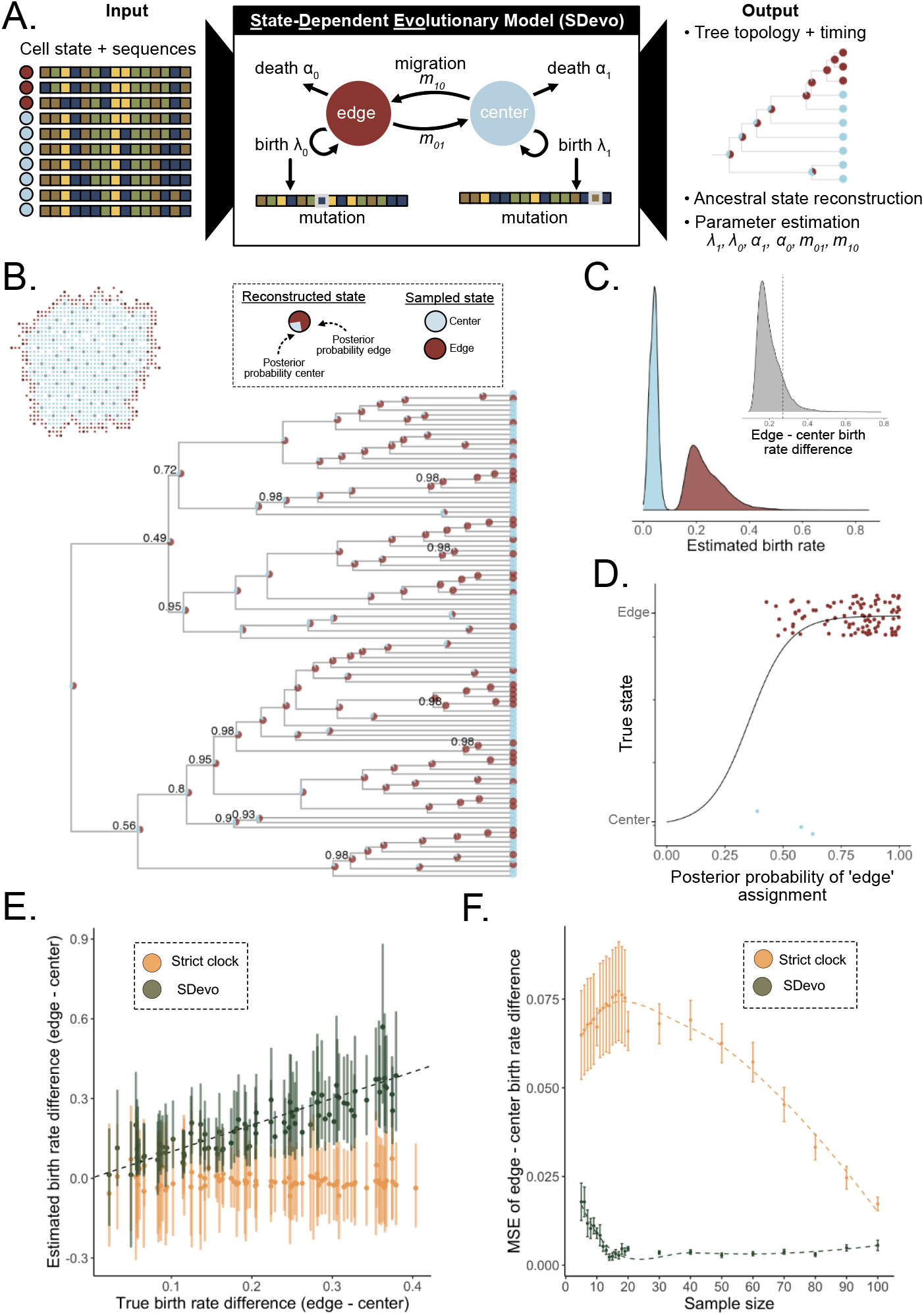
State-Dependent sequence evolution model (SDevo) estimates boundary-driven growth in simulated tumors. **A**. SDevo model schematic. Using input sequences and states (edge in maroon versus center in blue) of sampled tumor cells, SDevo reconstructs a tree with ancestral states (state probabilities represented by node pie charts) and estimates model parameters. SDevo links state-dependent clock rates to birth rates. **B**. Reconstructed time tree estimated by SDevo on an example simulated tumor (α = 0.145). At each internal node, the posterior probabilities for ancestral edge or center states are shown as a pie chart. Clade posterior support is indicated if less than 99%. The inset shows the sampling scheme for the tumor. **C**. Marginal posterior distributions of estimated edge and center birth rates, which are summarized by birth rate differences between edge and center cells (inset, dashed line indicates the true difference). **D**. Posterior probabilities of ancestral state reconstructions versus true state assignments. **E**. SDevo (green) estimates of birth rate differences between edge and center samples across a variety of true birth rate differences (α varies between 0 and 0.435, *n* = 50) compared with estimates under a strict clock (gold). Points and bars represent mean and 95% HPD intervals, respectively. Dashed line is *y* = *x*. **F**. Mean squared error (MSE) of estimated birth rate differences in simulatedtumors (α varies between 0 and 0.435) versus input sample size for SDevo (green) and strict clock sequence evolution (gold) models. Error bars represent standard error of MSE.

We first demonstrate the utility of SDevo on simulated tumors undergoing boundary-driven growth. From the genetic sequences and labeled cell-states for sampled cells isolated at a simulated tumor endpoint (Figure 3B inset), SDevo reconstructs the most likely relationship among sampled cells and the time at which those cells diverged (Figure 3B). The birth rates for edge and center-associated cells are inferred from the branching and mutational structure of sampled extant cells (leaves on the tree), permitting quantification of overall birth rate differences between the two spatial compartments (Figure 3C). SDevo correctly identifies that boundary-linked cells have a higher birth rate than center-linked cells (mean edge birth rate advantage = 0.20, 90% HPD = 0.12 - 0.29, true value = 0.27 in the representative simulation). SDevo additionally reconstructs the probability of each spatial state (center versus edge) for the ancestors of the sampled population (plotted as pie charts on the internal nodes of Figure 3B). These reconstructions suggest that the majority of ancestors divided on the tumor edge, consistent with the findings of ***Househam et al. (2022***) and our expectations of boundary-driven growth. SDevo’s Bayesian approach further quantifies confidence in its ancestral reconstructions: ancestral cells with the highest posterior probability of existing on the tumor edge were indeed likely to have divided there (Figure 3D). On the other hand, cells with more uncertain ancestral reconstructions are less likely to have been on the tumor edge at division (Figure 3D). Finally, we applied SDevo to tumors simulated under a range of spatial constraints (see Materials and Methods). We find that at a moderate sample size (*n* = 50), SDevo is able to accurately quantify birth rate differences, whereas a two-state birth-death model without a state-dependent clock (BDMM’ under a strict clock) fails (Figure 1E). We further observed that SDevo remains accurate for as few as 10 samples, whereas a strict clock model requires >100 samples to reach close to the same accuracy (Figure 1F).

### SDevo is robust to a variety of sampling approaches and tumor growth modes

To evaluate SDevo’s strengths and limitations in clinical tumors, we sought to validate that SDevo detects boundary-driven growth under various sampling strategies. Whereas in the initial simulation studies we maximized the distance between sampled cells (i.e. diversified sampling), we also implemented a random sampling scheme as might be present in single-cell studies (Figure S4A). Under random sampling, cells sampled close together provide minimal additional genetic information, but may create spurious signatures of rapid branching. Despite this, SDevo successfully estimates edge-driven birth advantages from randomly sampled cells (Figure 4A). In contrast, even with a large number of cells sampled (*n* = 100), the strict clock multi-type birth death model often fails to detect the same birth rate differences (Figure S4B). We also assessed SDevo’s robustness to punch biopsy sampling, in which a population of nearby cells are captured. We biopsy-sampled our simulated tumors, and only called mutations exceeding a 0.3 cellular fraction threshold within a punch (see Materials and Methods). We find that while punch-style sampling adds more random error due to variation in sampled diversity, especially in tumors with high turnover rates, SDevo largely still detects state-dependent birth rate effects (Figure 4B).

**Figure 4.**
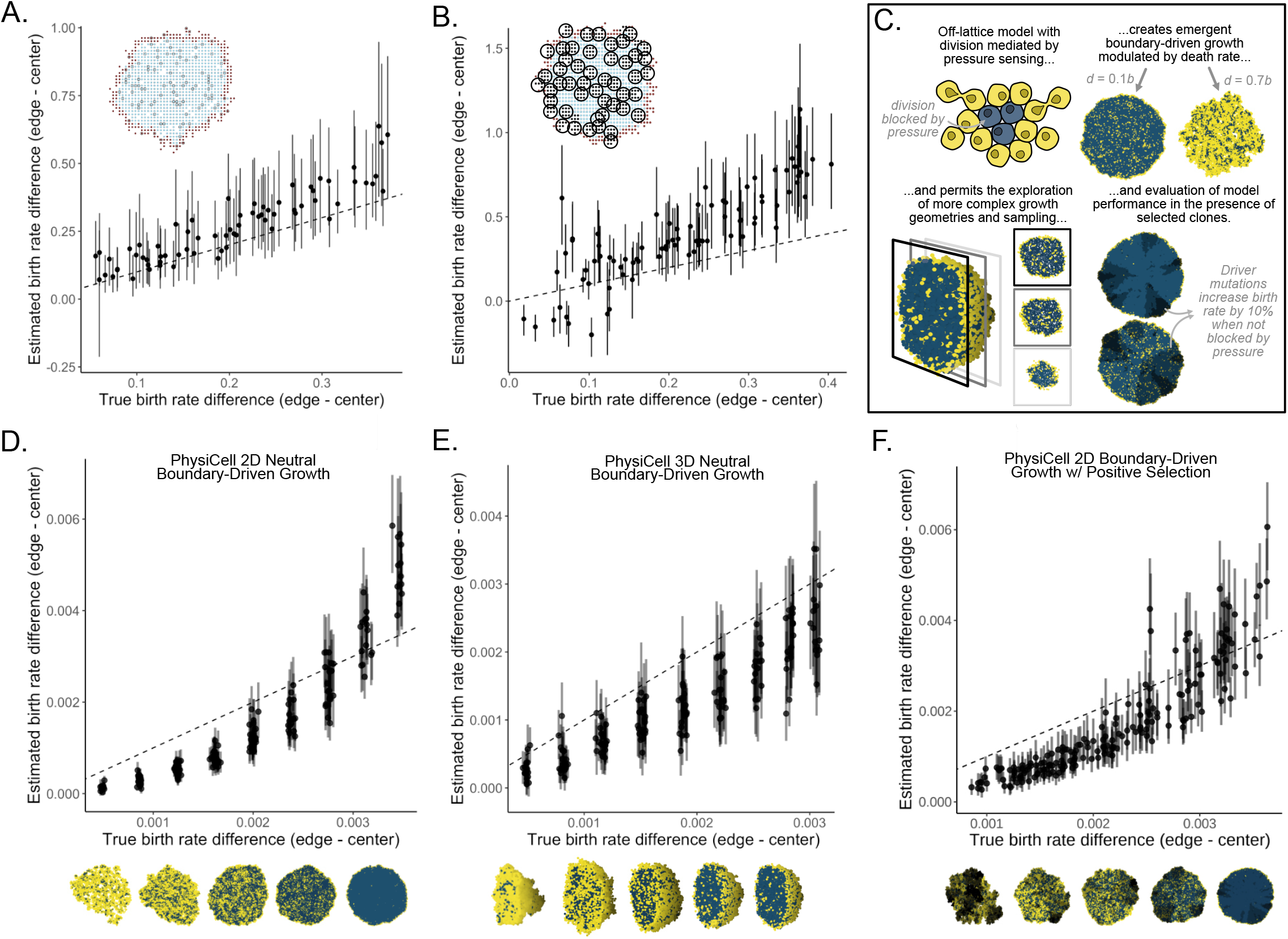
SDevo is robust to a variety of sampling approaches and growth modes. **A**. Estimated versus true edge - center birth rate differences with random sampling (*n* = 100 cells/tumor). **B**. Mean and 95% HPD intervals of SDevo estimates of birth rate differences between edge and center when sequences are constructed from variants above 30% frequency in a simulated punch biopsy (inset). **C**. Schematic of continuous-space tumor growth simulation governed by biomechanics using PhysiCell (***Ghaffarizadeh et al., 2018***). Only cells under low physical mechanical pressure from their neighbors (visualized in yellow as opposed to blue) can divide, generating boundary-driven growth. SDevo recovers growth rate differences generated through variable death rates under **D**. neutral 2D growth, **E**. neutral 3D growth, and **F**. 2D growth in the presence of strong driver mutations (μ_*driver*_ = 0.01, *s*_*driver*_ = 1.1, see Materials and Methods). Tumor snapshots below the x-axes show representative examples of growth dynamics under variable death rates (right to left: *d* = 0, 0.2, 0.4, 0.6, 0.8) and resultant pressure. In the case of the tumors under selection shown in **F**., darker colors represent cells with driver mutations.

Next, we assessed SDevo’s robustness to more complex growth by exploring an off-lattice model, a more flexible class of spatial models also employed to study tumor evolutionary dynamics (***Rejniak and Anderson, 2011***; ***Jeon et al., 2010***; ***Ozik et al., 2018***). We simulated under a continuous space model of tumor growth implemented using the agent-based cellular engine PhysiCell (***Ghaffarizadeh et al., 2018***). To mimic boundary-driven conditions, we linked division probability to mechanical pressure - cells crowded by their neighbors could not divide (see Materials and Methods, Figure 4C). As in the lattice-based simulations, higher cell turnover relaxes mechanical pressure, modulating spatial constraint. We first verified that SDevo continued to identify birth rate differences in these more complex simulations. We simulated 2D neutral growth, and found that SDevo sensitively detects an elevated birth rate at the tumor edge, even when birth rate differences were minimal (Figure 4D). However, SDevo slightly underestimates the birth rate differences at high death rates (i.e., low birth rate differences). We also confirmed that SDevo was robust to spatial division rate heterogeneity induced by increasing cell migration, as opposed to cell death (Figure S8A), and to a sigmoidal pressure threshold for cell proliferation (Figure S8C, D). We next simulated tumors grown in 3 dimensions and sampled across multiple *z*-slices, mimicking clinical sampling approaches. We determined that SDevo accurately reconstructs birth rate differences, albeit with wider posterior intervals (Figure 4E). We note that trees reconstructed from the 3D simulations tend to deviate more from expected edge-biased branching patterns than those from the 2D simulations (Figure S5), reflecting more complicated growth dynamics and potential obfuscation via the sampling scheme. These observations further highlight the necessity of incorporating both branching and clock rate patterns to quantify boundary-driven growth in clinical scenarios.

Finally, we tested the extent to which SDevo detects boundary-driven growth dynamics when both spatially-determined and cell-intrinsic fitness differences influence growth, as the action of strong positive selection has been previously shown to distort the shape of tumor phylogenetic trees (***Chkhaidze et al., 2019***; ***Househam et al., 2022***; ***Li et al., 2021***). We find that SDevo continues to detect differences in birth rates between center and periphery-associated cells even in the presence of strong selection (Figure 4F and S8B, see Materials and Methods). Notably, even as lineages with driver mutations expand, these cells are still subject to spatial constraints. As a result, similar patterns of branching and clock rate differences between center and periphery-associated cells re-emerge. However, we anticipate that if cell death is sufficiently high, a driver mutation could lead to rapid expansion of a center-bound lineage and mask signals of boundary-driven growth.

### SDevo detects growth rate differences in hepatocellular carcinomas

To quantify boundary-driven growth in a clinical tumor setting, we applied SDevo to multi-region sequencing data of two hepatacellular carcinoma (HCC) cancers published by ***Li et al. (2021)***(Figure 5). The authors sequenced two HCC tumors from a single patient, carried out 3-dimensional spatial micro-biopsy sampling followed by whole-genome sequencing (Figure 5A and E) and classified punches as “edge” or “center”. The genetic maximum likelihood trees of each tumor (Figures 5B and F) qualitatively demonstrate an increased genetic divergence at edge punches. To apply SDevo, we created input pseudo-sequences for each punch using three independent 25,000 SNV random subsets of those identified in the original study. We assumed unidirectional transition from edge to center, in line with biological expectations of solid tumor growth, to constrain death and transition rate parameter space (see Materials and Methods). SDevo jointly reconstructed tumor time trees along with the most likely ancestral internal node states. From these results we infer that while most ancestral cells divided on the tumor periphery, some population expansion occurred in the tumor center. We note that we do not use a predefined outgroup for this analysis, so there is slight differences in rooting for these time trees compared to the genetic maximum likelihood trees. SDevo found strong support for birth rate differences between edge and center in both tumors (Figure 5D and H). We estimated that cells on the edge have a mean 6.05x birth rate advantage over center cells in Tumor 1 (95% HPD = 4.43 - 7.73x), and a mean 3.75x birth rate advantage in Tumor 2 (95% HPD = 3.35 - 4.16x) summarized across all SNV subsets. To assess how sensitive these results were to differences in state classifications or punch heterogeneity, we also called alternate edge/center states based on a threshold of 10% of the tumor diameter (∼2mm and ∼1.5mm for Tumor 1 and Tumor 2, respectively) from the schematic boundary (Figure S6A and E). We found consistent results for Tumor 2, but observed that Tumor 1’s alternate edge/center classifications showed more variable and reduced support for boundary-driven growth, which was not unexpected given that the alternate states updated the classification of previously center-assigned punches with less genetic divergence to edge (Figure S6D and H). We further found consistent results when removing a single punch from Tumor 1 (Figure S10), which may have captured multiple subclones (Figure S9).

**Figure 5.**
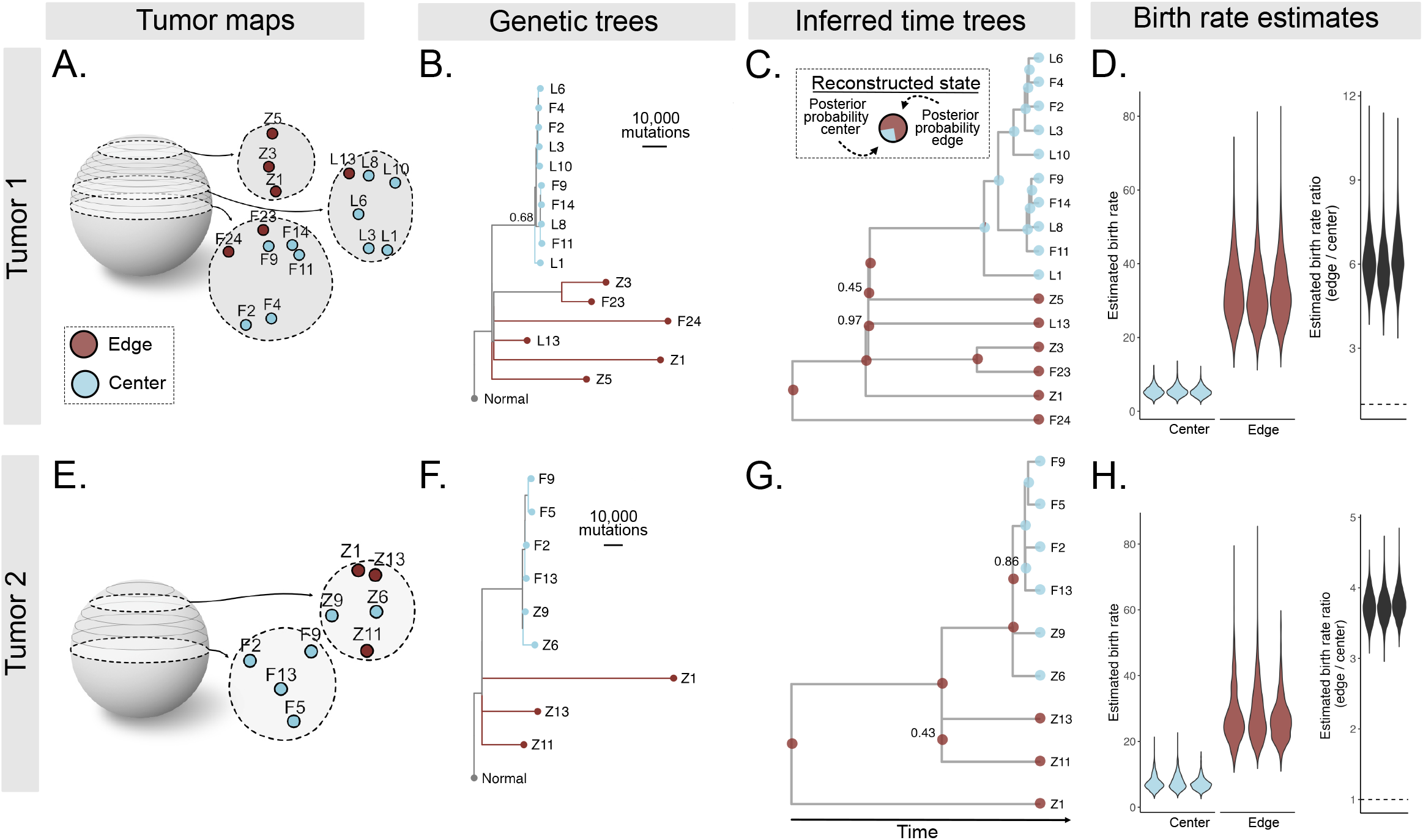
Quantification of boundary-driven growth in hepatocellular carcinomas. **A**. Multi-region 3D sampling map for Tumor 1 adapted from ***Li et al. (2021)***. Sampling locations are marked and labeled in *z*-slices, and center and edge classifications (taken from the original study) are shown in blue and maroon, respectively. **B**. Maximum likelihood genetic tree reconstructed from all variable sites. Node confidence is labeled if less than 99%. **C**. Tumor 1 phylogeny reconstructed from pseudo-sequences of one subset of variable sites. Tip colors indicate sampled punch state and pie charts on internal tree nodes represent posterior probabilities of ancestral state reconstructions. Clade supports are indicated at nodes if less than 99%. **D**. Marginal posterior distributions for edge (maroon) and center (blue) birth rates across three independent subsamples of variable sites, and estimated edge to center birth ratio (mean 6.05x). Dashed line marks ratio of 1. The same plots are shown for an additional HCC tumor (Tumor 2) from the same dataset (**E-H**), estimating a mean 3.75x edge to center birth rate ratio. Tumor schematics are vertically inverted for ease of visualization.

Although we inferred a higher birth rate on the edge in these clinical tumors, the branching rate patterns in Tumors 1 and 2 qualitatively did not match our expectations from simulations. These branching patterns are potentially influenced by selection, as noted originally by ***Li et al. (2021)***, or by the non-uniform sampling scheme (Figures 5A and E). Likely due to these branching patterns, we find a strict clock model, which assumes independence of sequence evolution and cell division, did not detect boundary-driven growth. Instead it estimated that center cells have a slightly higher birth rate (Figure S7). We note that the sample sizes of Tumor 1 and Tumor 2 were well below the sample size requirements in simulations to detect boundary-driven growth with a strict clock model (Figure 3F). In addition, we found that incorporating a state-dependent sequence evolution model changed the estimated internal node timings (Figure S7C and F). Specifically, reconstructed center-bound nodes were estimated to have occurred more recently under a strict clock than under a state-dependent evolution model in which center cells would be expected to divide less frequently.

## Discussion

Tumor evolutionary progression is a complex process driven by genetic, epigenetic, environmental, and immune factors. Quantitatively disentangling the contribution of spatial factors to tumor growth dynamics is an important component of both reconstructing tumor clinical histories and predicting future growth. Our understanding of spatial drivers of tumor growth has largely been informed by xenograft models, as we have had limited ability to assay for these effects in clinical tumors. Here, we introduce SDevo, a new Bayesian phylodynamic model that learns differential cell birth rates of discrete classes (here, tumor periphery or center-associated). Although SDevo is general in scope and applicability, in this study, we demonstrate that it successfully infers birth rate differences between the tumor edge and center from multi-region sequencing data. We show that SDevo is relatively robust to sampling choices (i.e., punch biopsies and locations) and biological factors (i.e., cancer driver mutations and 3D versus 2D growth modes). We further find quantitative evidence for boundary-driven growth in clinically-derived hepatocellular carcinomas resected at a single time point.

Our assessment of boundary-driven growth in HCC quantitatively expands the observations of ***Li et al. (2021)***. The authors originally hypothesized that Tumor 1’s tree structure matched a simulated scenario of boundary-driven growth followed by the expansion of a selected clone in the center, and that Tumor 2’s tree structure matched dominant boundary driven growth. The authors made these assessments by simulating tumors and comparing the distributions of clones and variant allele frequencies to the sequenced tumors. They further noted that genetic divergence was higher in punches collected from the tumor periphery.

Our study quantifies these patterns by estimating these birth rate differences directly with joint inference of tree topology and sequence evolution. Notably, although small sample sizes, clustered sampling, and the hypothesized selection for an internal clone in Tumor 1 may have distorted the branching structure of the trees, SDevo is able to detect past boundary-driven growth from clock rate differences. By explicitly incorporating the mutational process, SDevo leverages data more effectively than models that only learn from state-dependent branching. This approach is particularly important when only a few areas of a tumor are sequenced. These findings, along with previous *in silico* evidence that selection changes the shapes of tumor trees (***Chkhaidze et al., 2019***; ***Yang et al., 2022***), highlight the importance of employing multiple tree patterns to quantify interacting modes of tumor growth. Although future work should more comprehensively profile how multiple spatial and non-spatial drivers of growth can impact observed tree patterns, our analysis of non-neutral tumors (Figures 4F and S8B) suggests that SDevo can detect boundary-driven growth in the presence of selection.

Quantifying the impact of spatial restrictions on clinical tumor growth informs how we understand, predict, and control cancer evolution. A robust literature has established that boundary-driven growth modulates the efficiency of positive and purifying selection (***Kayser et al., 2019***; ***Fu et al., 2022***), alters overall growth rates (***van der Heijden et al., 2019***; ***Colom et al., 2020***), and increases the efficacy of adaptive therapy (***Bacevic et al., 2017***; ***Strobl et al., 2022***; ***Fusco et al., 2016***). Spatial restrictions also change the expected distribution of genetic variation in solid tumors (***Sun et al., 2017***; ***Ahmed and Gravel, 2018***; ***Waclaw et al., 2015***; ***Fu et al., 2022***), and impact how clinically informative biopsies should be collected (***Kostadinov et al., 2016***). Although we find robust evidence for boundary-driven growth in HCC, its prevalence and strength likely varies by stage of tumor growth and tumor type (***Noble et al., 2022***). For example, increased vascularization, cellular migration, physical anatomical structures, or tumors reaching a local carrying capacity could alter spatial growth restrictions. Further applications of SDevo to other tumor cases and types will enable us to explore nuances of these growth phenomena.

Importantly, the utility of SDevo is not limited to understanding the impact of boundary-driven growth, but in fact, can be applied in any instance in which sequenced tumor samples can be classified into discrete, observable states. Immediately, SDevo could be extended to test other proposed tumor growth modes – for example, growth against a solid surface, such as bone in osteosarcoma, along a unidirectional invasive front (***Ryser et al., 2020***), or in different glandular compartments (***West et al., 2021***). Because tumors can grow under a wide variety of anatomical constraints, integrating system-specific factors can help assign biologically-relevant environmental states for the application of SDevo (i.e., edge categorization may constitute those cells that have penetrated the basal layer as opposed to those that are most radially extreme). Even more broadly, SDevo could be applied to study the growth impacts of other environmental or cell-intrinsic factors, for instance, immune invasion, hypoxia, metastatic versus primary sites, or genetic features, by decomposing complex phenotypes into discrete states.

Phylodynamic approaches such as SDevo have major advantages compared to our current approaches for estimating evolutionary information from tumors, namely approximate Bayesian computation (ABC) (***Beaumont et al., 2002***) or other approaches that compare simulated and clinical tumors via summary statistics (***Noble et al., 2022***). To be clear, these approaches have yielded extensive insights into tumor evolution, including patterns under boundary-driven growth (***Sottoriva et al., 2015***; ***Sun et al., 2017***; ***Chkhaidze et al., 2019***; ***Househam et al., 2022***). However, these approaches are computationally costly, requiring the generation of often tens or hundreds of thousands of simulated tumors, on which one must compute extensive summary statistics. In addition, ABC comes with technical challenges, including the necessary choice (and potential unavailability of) low dimensional sufficient summary statistics. Although Bayesian phylodynamics comes with its own technical challenges (i.e. identifiability, sensitivity to model assumptions, choice of priors, see ***Louca and Pennell (2020)***; ***Louca et al. (2021)***), it does not require tumor simulation. Furthermore, the generality of discrete traits affecting growth dynamics means it is easily adaptable to answer new questions. While both ABC and phylodynamics offer ways to understand clinically-derived samples, the full promise of phylodynamics has yet to be widely exploited.

Phylodynamic approaches to understanding tumor evolution offer additional benefits: 1) used in conjunction with well-calibrated molecular clocks, inferred trees can help estimate the timing of clinically-important events, such as the emergence of subclones or metastatic events. While these analyses have been employed in the context of uniform growth rates (***Lote et al., 2017***; ***Hu et al., 2020***; ***Alves et al., 2019***), the expansion of tree models to permit differential birth rates could improve timing accuracy. 2) Incorporating differential growth rates across a tree can lead to more accurate tree topologies, as has been demonstrated in influenza evolving in multiple host species (***Worobey et al., 2014***). 3) Inferring ancestral states can elucidate population history and tumor evolutionary processes at time points that cannot be clinically sampled. Recently, ***Zhao et al. (2021)*** analyzed the intra-tumor spatial and genetic architecture of renal cancers and concluded cells in the tumor center are more likely to seed metastasis. However, the study was limited to observing the *extant* position of these samples, whereas SDevo reconstructs these states at the time of clinical events (i.e. divergence of a metastatic clone). These three points suggest more broadly how tumor trees can be leveraged to gain new quantitative insights into tumor evolution, and demonstrate the broad utility of modeling evolutionary processes on trees.

Beyond its application to cancer evolution, SDevo is a novel phylodynamic model with broad usefulness to incorporate state-dependent clock rates into evolutionary inference. While the field of phylogenetics has developed a broad array of clock models, SDevo represents the first model in which clock rate is linked to population birth. SDevo could be particularly useful in microbial and viral populations where diversification and mutational accumulation operate on similar timescales, and may be linked to underlying state variables (for example, location). We demonstrated that incorporating clock rate differences, instead of relying solely on tree diversification rates (as in BDMM’ and other multi-state birth-death models (***Maddison et al., 2007***; ***Kühnert et al., 2016***; ***Vaughan, 2022***)), can improve inference in cases where sampling may be non-uniform. This may be particularly important when sampling rates vary - for example, countries with variable rates of molecular surveillance for SARS-CoV-2. To facilitate access to these methods, SDevo will be released as a package in the popular Bayesian phylogenetic platform BEAST2 (***Bouckaert et al., 2019***) and is available on GitHub before publication. As with all phylodynamic models, identifiability represents a pervasive concern, but incorporating biological knowledge for determining priors can help constrain the model space. In our analysis of HCCs, we use information about cell transition and death rates to distinguish between multiple parameters that impact trees and estimation in interrelated ways.

Biological complexity within tumors can complicate SDevo’s application and interpretation via spatiallyor temporally-varying selection. First, strong selection can destroy or alter signals of boundary-driven growth (***Chkhaidze et al., 2019***; ***Li et al., 2021***). For example, a hard bottleneck, as in the cases of surgery or chemotherapy, would likely temporarily destroy signals of boundary-driven growth. Such signals would likely also re-emerge were the tumor to regrow via boundary-driven growth. Second, gain of driver mutations will lead to cell-intrinsic fitness differences that may not correlate with spatial location. Third, disentangling boundary-driven dynamics from other environmental or cell-intrinsic factors could be especially difficult under time-varying selection. For example, angiogenesis could increase resources to center cells later in tumor growth (***Junttila and de Sauvage, 2013***) and complex cell-to-cell interactions may create frequency dependencies that further complicate observed spatial patterns (***Karras et al., 2022***; ***Strobl et al., 2022***; ***Farrokhian et al., 2022***). We have shown that SDevo can detect signals of boundary-driven growth even with driver-induced selection, but future work should further probe this robustness.

Although SDevo is a powerful tool, we note several important limitations that require further caution when applying it to data. First, SDevo assumes mutations occur at cell division. If instead, most mutations emerge due to exogenous processes (***Abascal et al., 2021***), birth-driven genetic divergence could be masked. While this might decrease SDevo’s power, exogenous mutational processes distributed evenly across a tumor are unlikely to generate false positive signals of boundary-driven growth. Second, extensive cell mobility could weaken signatures of boundary-driven growth even if boundary-associated cells have birth rate advantages. Third, as we demonstrate in Figure 3, sample sizes must be sufficient to detect state-dependent effects. We maximize limited sample sizes by choosing priors that are biologically informed (for example, unidirectional state transitions), but larger sample sizes will enable inference with less informative priors. Data sets that meet this requirement are becoming rapidly available, so we anticipate phylodynamic models such as SDevo becoming increasingly powerful.

The expanded application of phylodynamics to cancer sequencing data relies both on developing methods to exploit single-cell sequencing data (***Chen et al., 2021***; ***Moravec et al., 2022***), and understanding the relationship between sequenced multi-region punches and the many single cells that comprise them. As has been noted previously, multi-region sequence trees are not phylogenies (***Alves et al., 2017***), and punch-wide genetic composition does not necessarily capture all cellular genotypes (***Caravagna et al., 2020***). Although SDevo is fairly robust to our simulated punch-style sampling and we analysed HCC data from small, largely homogeneous punch biopsies, best practices for applying phylodynamic models to trees of deconvoluted clones is an important area for future research.

Applying phylodynamic methods to tumor populations is in its infancy, but new methods that overcome the barriers of working with tumor data will help extend the applicability of these approaches (***Alves and Posada, 2018***; ***Chen et al., 2021***). Here, we demonstrate the utility of phylodynamic models in quantifying spatial factors driving cancer progression. As technologies enabling the widespread and high-throughput generation of tumor trees advance (***Yang et al., 2022***; ***Lim et al., 2020***), we expect adapted phylodynamic approaches such as SDevo to provide a rigorous analytical toolkit for extracting quantitative insights from these data.

## Materials & Methods

### Tumor simulations

#### Eden model

An agent-based model was implemented in Python3 which places simulated cells on a 2D lattice. Simulations are initiated with a single cell in the center of the lattice. Cells attempt division with rate λ (per day) or die with rate α (per day). At each time step Δ*t* (1/24 days, or 1 hour), these rates translate to probability of division λΔ*t* and probability of dying αΔ*t*, which is a suitable approximation when the product of the rate and the time step is much less than 1. Under boundary-driven growth, cells only successfully divide if there is an empty lattice spot in its Moore neighborhood. If multiple neighboring spaces are available then the cell randomly chooses the location for its daughter cell from open neighboring spaces. Under unrestricted growth, if a cell attempts division, its daughter cell will occupy an empty lattice spot in the Moore neighborhood if available, but if not, the cell will still divide and push cells in a random direction to make space. Overlapping cells are pushed in the same direction until a neighboring lattice spot is available, which the pushed cell will occupy. In both simulations, if a cell divides, each daughter cell can gain mutations with probability μ (per division). Mutations are then drawn from a Jukes-Cantor model of sequence evolution and follow an infinite-sites assumption. Therefore, each time a mutation is gained, a site is added to all cells in the simulation. Simulations are stopped when the number of living cells is more than 1000. The ground truth birth rates are assessed at discrete time points in the simulation by recording the current state of each cell and the proportion of cells that have progeny in the next time step. True birth rates are considered to be the mean across all time steps weighted by the number of cells in each category. This method calculates effective birth and death rates on the edge and center given the simulated spatial constraints by calculating empirical division rates on the edge and center of cells through simulated time. Effective spatial constraints in the boundary-driven model were controlled by changing cell death rate, where increased cell turnover allows center-trapped cells to divide more readily (Figure S1). Input parameters are proliferation and death scalars, where λ = (1 − *death*)(*proliferation*)/2 and α = *death*)/2. To evaluate the accuracy of parameter estimation, we ran 1000-cell tumor simulations with *proliferation* = 1, μ = 1 and a range of *death* = (0 − 0.87). While clinical tumors have large variability in rates of proliferation, death, and mutation, these parameters fit within this biological range (***Bozic et al., 2010***; ***McFarland et al., 2013***; ***Tomasetti et al., 2013***; ***Waclaw et al., 2015***).

#### Eden tree statistics

Tree statistics in Figures 1 and 2 were calculated from simulated tumor trees that include all extant cells. Normalized terminal branch lengths were calculated by dividing terminal branch lengths of tumor time trees by total simulation time. Clock rates were calculated by dividing total number of mutations accumulated in each alive cell by simulation time. Edge and center states for terminal branch lengths are defined by cell location at the end of the simulation, where edge cells are defined by being the most extreme cell on either the X or Y spatial axis for each row and column, respectively, or within one cell of this boundary. Fraction of the lineage time spent on the edge is determined by averaging across all lineage node states weighted by time tree branch lengths.

#### Continuous space model

To probe the robustness of SDevo to more complex selective events and higher dimensions, we implemented an additional set of simulations in the physics-based cellular simulator, PhysiCell (***Ghaffarizadeh et al., 2018***). Briefly, PhysiCell is an open-source, agent-based model implemented in C++ in which cell movement is governed by biomechanical interactions among cells. To simulate boundary-driven growth, we created a PhysiCell instance in which cells are only able to divide when under low mechanical pressure, using the cell state variable, *simple_pressure*. As a result, similar to the Eden model, most cell division is restricted to the tumor periphery, or to cells with adjacent space created by the recent death of a neighboring cell. Cells initially divide at a rate we arbitrarily set to 1, except when above the pressure threshold, τ, in which case, they divide at rate 0. We also explored a sigmoidal relationship between pressure and birth, where the birth rate *b* = 1 − (1 + *exp*(−5 ∗ (*pressure* − τ)))^−1^. Cells die at rate *d*, regardless of their pressure status. To simulate selection, during each cell division, a daughter cell can acquire a driver mutation conferring a 10% fitness advantage (***Beerenwinkel et al., 2007***) with probability μ_*driver*_, which acts multiplicatively (i.e. a cell with two drivers has a 21% faster growth rate than one with 0) (***Weile et al., 2017***). Tumors are grown to a final size of *N* extant cells, of which *n* are sampled. After the simulation, a Poisson-distributed number of neutral mutations is augmented to each cell division with λ = μ_*passenger*_. Using the continuous space model, we investigated all pairwise combinations of 2D and 3D, neutral and selective scenarios, and ran 25 tumor simulations for each combination of parameters (τ = 1, *d* = (0, 0.1, 0.2, 0.8), μ_*pass*_ = 1, *n* = 100), except for 3D selection, where we simulated *d* = (0, 0.2, 0.6, 0.8) with 10 tumors each. For the 2D models, *N* = 10, 000 and for the 3D model, *N* = 15, 000. For the selective model, μ_*driver*_ = 0.01 and for the neutral models, μ_*driver*_ = 0. Note, we used a value of μ_*driver*_ well above expected rates of driver mutations (≈ 10^−5^, (***Bozic et al., 2010***)) to conservatively test SDevo in an extreme case of selection. To probe SDevo’s performance when cellular constraint is reduced by migration instead of cell death, we performed 10 simulations at *d* = 0.2 where cells migrate according to unbiased Brownian motion at 0, 0.5, 1, 1.5 or 2μ*m*/*minute* (all other parameters as above). To probe SDevo’s robustness under a sigmoidal relationship between pressure and birth rate, we ran 10 simulations with *d* = (0, 0.2, 0.4, 0.6), and all other parameters as above. One outlier in the 3D boundary-driven growth simulations was removed due to convergence on a local optimum. Ground truth edge and center birth rates were determined by first classifying cells as within 10 microns (approximately 1 cell width) of the tumor periphery as edge, and those more than 10 microns from the edge as center. The average birth rate was computed separately within each of those classes over multiple discrete time points (10-40, depending on the overall rate of tumor growth) and combined by a weighted average according to the number of cells at each time point. Cells under too much pressure to divide at the sampled time (*simple_pressure* > τ) were calculated as having an instantaneous birth rate of 0.

#### Sampling procedures

2D simulations were sampled by maximizing the distance between sampled single cells in physical space (diversified sampling). This ensures that a sufficient number of edge and center classified cells were sampled and that sampled cells were not clustered. Bulk punch biopsy sampling was mimicked by choosing a center cell and a target of 8 cells immediately surrounding that were grouped into a single punch. Punches were iteratively drawn and shifted if they overlapped with a previously punched group of cells. Sampling ended when the target number of punches was reached (50 punches) or sampling was no longer possible without significant overlap. Punch sequences were generated using all mutations above a cellular fraction cutoff of 0.3. 3D sampling was approximated by taking 5 simulated slices through the tumor *z*-plane at 2/8ths, 3/8ths, 4/8ths, 5/8ths and 6/8ths of the range of the *z* values of a given tumor. Within each slice, cells were sampled to maximize the inter-cell distance, as described above, and the number of cells per slice was proportional to the number of cells in the slice relative to the number of cells across all slices.

#### Multi-type birth-death models applied to boundary-driven growth

The birth-death process describes how lineages duplicate (birth), die (death), and are sampled (where samples are tips on a phylogenetic tree) (***Gernhard, 2008***). The multi-type birth-death model extends this by considering birth, death and sampling to occur in different states (sometimes also referred to as different sub-populations, traits, or types) and how lineages jump between these states. The rates of birth, death and sampling vary depending on the state of a lineage. For the case of boundary-driven growth, we model a two-state process, with one state denoting cells in the center of the tumor and the other state denoting cells on the edge of the tumor.

#### Posterior Probability

To perform Bayesian inference, we define the posterior probability *P* (*T, σ, θ* |*D*) of the timed phylogenetic tree*T*, the evolutionary model and parameters (*σ*), and the population model and parameters *θ*, given the data, *D*. This posterior probability is typically expressed as:

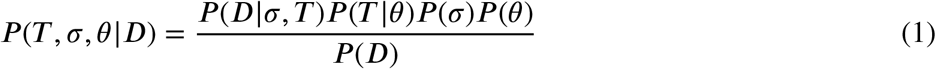

In the case of the state-dependent multi-type birth-death model, we cannot assume the tree likelihood (*D*|*σ, T*) and the tree prior *P* (*T* |*θ*) to be independent, as the rate of evolution directly depends on the population model. In other words, how fast evolution happens on a lineage depends directly on the state of that lineage. We therefore define ℋ as a mapped state transition history, that contains a random mapping of state change events given a set of parameters *θ* of the multi-type birth-death model. We then define the tree likelihood as *P* (*D*|*σ, θ, T*, ℋ). Additionally, we say that instead of computing *P* (*T* |*θ*) directly, we only compute the tree prior for one realization of the state transition history, i.e. *P* (*T*, ℋ |*θ*). The posterior probability then becomes:

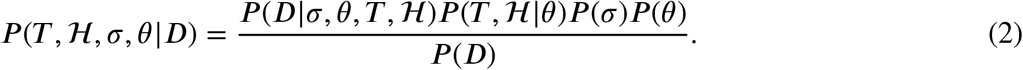

Performing MCMC inference to characterize this posterior probability distribution would require integrating over all transition histories ℋ using MCMC. This is overall incredibly slow and limits the application of the method. Instead, we formally integrate over all possible histories ℋ, to get the following posterior probability:

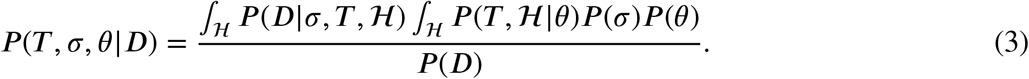

*P* (*T*| *θ*) = *∫*_*H*_ *P* (*T*, ℋ |*θ*) is computed as described in ***Kühnert et al***. (***2016***), which is achieved by treating the states of lineages probabilistically instead of discretely.

Lastly, we set *∫* ℋ *P* (*D*| *σ, T*, ℋ) = *E*[*P* (*D* |*σ, θ, T*, ℋ) = *P* (*D*| *σ, θ, T, E*[ℋ]), with *E*[ℋ] being the expected/average state transition history, which contains, for each lineage *i* in the phylogeny, its expected time spent each state *s*. This leaves us with:

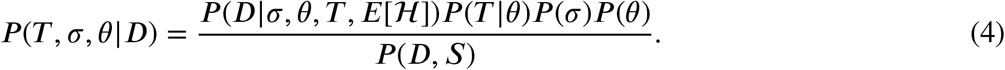

#### Modeling birth-dependent evolution

In order to model different rates of evolution for different states, we first compute the expected time each lineage in the phylogenetic tree *T* spent in each state. To do so, we use a stochastic mapping approach related to those described in ***Nielsen (2002)***; ***Huelsenbeck et al. (2003)***. We first compute the probability 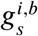 of each lineage *i* in the phylogenetic tree being in any possible state *s* over time *t* from the tips to the root as described in ***Kühnert et al. (2016)***. These state probabilities are conditional only on events that occurred more recently than *t* and therefore not on all events in the phylogeny. During this backwards propagation, we keep track of the time dependent transition matrix *Q*(*t*)^*i*^ that describes the rate of probability flow between any two states at time *t* due to state transitions or birth events between states. As a result, once we reach the root, 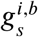contains all events in the phylogeny and is therefore equal to 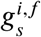, i.e. the forward probability of lineage *i* being in state *s*.

Following ***Stolz et al. (2022)***, we first define 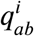 as:

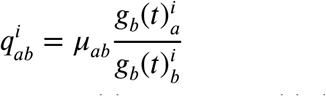

with *μ*_*ab*_ being the rate of state change due to state transitions or cross-birth events.

We then compute the probabilities of any lineage being in any possible state conditional on all events in the phylogeny 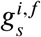 forwards in time as:

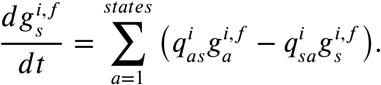

By keeping track of the forward probabilities 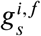 on each lineage, we can then compute the expected time 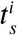 that lineage *i* spends in any of the possible states *s*. The values for 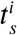 make up the entry for *E*[ℋ] in the posterior distribution (4). We then say that *c*_*s*_ is the rate of evolution, that is the clock rate, of a lineage in state *s*. Next, we compute the average rate of evolution on branch *i, c*^*i*^ as

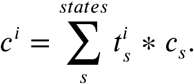

At each replication, error in copying the genetic material of a cell can occur. These errors tend to be more likely in cancer cells, where cellular control mechanisms are often faulty. Phylogenetic methods typically assume the evolutionary processes to be independent of population processes, such as cell replication. To model mutations happening at birth events, we assume that the birth rate *b*_*s*_ in state *s* and the clock rate in state *s* are proportional such that *c*_1_ = *c*_*avg*_ *b*_1_, *c*_2_ = *c*_*avg*_ *b*_2_, …, *c*_*n*_ = *c*_*avg*_ *b*_*n*_.

#### Implementation

We implemented the multi-type birth-death model with state-dependent clock rates as an addition to the Bayesian phylogenetics software BEAST2. SDevo depends on BDMM-prime (https://github.com/tgvaughan/BDMM-Prime) to compute the tree prior *P* (*T*| *θ*). To model mutations occurring at cell division, we set the relative rate of evolution in the different compartments (edge and center) to be proportional to the birth rates in these compartments. The implementation itself does not explicitly require this assumption and the relative rates of evolution can also be treated as a distinct parameter in the inference.

#### Validation

To validate the implementation, we perform a well-calibrated simulation study. In it, we simulate phylogenetic trees under a two-state birth-death model in which we assume the rate of evolution to be proportional to the birth rate in either compartment. We randomly sample the birth, death, and transition rates from the prior distribution, while fixing the sampling rate to 0.001 and then simulate a phylogenetic tree using MASTER (***Vaughan and Drummond, 2013***). We then simulate genetic sequences on top of the phylogenetic trees using different rates of evolution depending on the lineage’s compartment. Next, we infer the birth, death, and transition rates from the genetic sequences and show that the 95% highest posterior density (HPD) interval cover the truth in 95% of the 100 runs (see Figure S2).

#### SDevo application to simulated tumors

We applied SDevo to outputs of the Eden and PhysiCell simulations generated as described above. For each simulated tumor we calculated clock rate (*mutations/tree length/sequence length*) and edge and center sampling rates (*sampled / alive cells*). We set exponential priors on birth, death, and transition rates. Full parameterization can be reproduced from XML templates. MCMC chains were run to convergence. We used only chains that had an minimum effective sample size (ESS) for birth rate parameters greater than 200 for analysis. We summarized the output posterior distributions by mean and 95% HPD intervals. We further inferred maximum clade credibility (MCC) trees with median heights using BEAST 2.6.2 TreeAnnotator (***Bouckaert et al., 2019***). TreeAnnotator also gives posterior state probabilities for each MCC internal node.

#### SDevo application to hepatocellular carcinoma tumors

To apply SDevo to the hepatocellular carcinoma data, we labeled punches based on edge/center state labels as published by ***Li et al. (2021)***, Table S8 (reproduced in Figures 5A and 5E). For alternate states (Figures S6A and S6E), we labeled punches as edge if they were located within approximately 10% (∼2mm for Tumor 1 and ∼1.5mm for Tumor 2) of the tumor diameter from the schematic boundaries. Slices were reported to be from tumor hemispheres. Assuming a 0.2mm slice thickness, we estimated that slices Tumor 1Z and Tumor 2Z fell within the boundary region. The original amplicon genotyping panel artificially increases the apparent diversity within some clones relative to others, so to avoid incorporating this bias into the model, we used only whole-genome sequenced punches. ***Li et al. (2021)*** identified a large number of SNVs (254,268 for Tumor 1 and 142,032 for Tumor 2). To reduce computational requirements and improve convergence, we generated input pseudo-sequences by randomly subsampling 25,000 variable sites. We summarized results across three independent subsamples for each tumor. We called presence or absence of a variant at each site based on a VAF cutoff of 0.05. Variant allele frequency histograms displayed single-peaked distributions characteristic of a single major clone per sample, with the exception of tumor sample T1L13 (Figure S9). To ensure Tumor 1 results were not driven by over-counting mutations across multiple subclones of T1L13, we repeated the analysis excluding this sample and found quantitatively similar results (Figure S10).

We use a GTR + Γ_4_ site model, a fixed clock rate of 0.3 (units are arbitrary as we only use sites which are variable relative to healthy cells), and estimate sampling proportion (uniform prior). We use log-normal priors for birth (mean=20, S=0.5) and death rates (mean=15, S=0.5), and edge to center transition rate (mean=5, S=0.5). Note that these units are also arbitrary and are not calibrated to clinical time. In applying SDevo to these tumors, we constrain the parameter space in several ways to adapt to having relatively few samples, only a single observed time point, and unknown sampling proportion. 1) We assume unidirectional transition so that cells can only move from edge to center but not vice versa. As we only have a few observed state transition events, the transition rates would otherwise be relatively poorly informed. 2) We set priors on mean birth, death, and transition rates across the two states. Birth and death priors are identical across both states, while transition rates priors are asymmetrical to inform unidirectional transition and enable convergence in a complex parameter space. Full parameterization can be found in the XML template. We combined posterior estimates across three independent runs for each tumor. We inferred MCC trees with ancestral state reconstructions with TreeAnnonator. In addition to the SDevo-inferred trees and parameters, we also generated maximum likelihood trees using FastTree (***Price et al., 2009***) and Augur (***Huddleston et al., 2021***) under a Jukes-Cantor model for each tumor using all reported variable sites. Homoplastic sites contributed to lower support for one node in the maximum likelihood tree of Tumor 1 (Figure 5B) and we masked homoplastic sites to enable convergence in Tumor 1 and Tumor 2 SDevo inferences. Homoplastic sites represented < 1% (Tumor 1) or 6% (Tumor 2) of all sites across all tumor samples. In Tumor 2, more than 2/3rds of homoplasies were between two edge-associated punches (Z1 and Z13) potentially pointing to subclonal mixing, which is supported by their proximal spatial locations. The remainder of homoplasies in Tumor 2 and all of the homoplasies in T1 were evenly distributed across punches. As a result, removal of homoplasies did not act to bias branch lengths across the tree, with the exception of T2Z1 and T2Z13. As these punches are on the edge of the tumor, this masking should a priori result in lower estimated birth rates on the edge and thus conservatively bias the results towards a more equal birth rate between edge and center.

## Code and data availability

Custom scripts were used for simulation studies and data analyses. All code and data to generate figures are publicly available. Scripts to replicate analyses and figures are available at github.com/blab/spatial-tumor-phylodynamics, including a local R package *tumortree* (github.com/blab/spatial-tumor-phylodynamics/tumortree), which can be installed to build trees from the simulation outputs. The source code for SDevo is here: https://github.com/nicfel/SDevo. The source code to run spatially-constrained PhysiCell simulations and generate trees can be found here: github.com/federlab/PhysiCellTrees. All other packages used for analysis and visualization are also open source (***Yu et al., 2017***; ***Wang et al., 2019***).

## Acknowledgements

The authors would like to thank Xuemei Lu for answering questions and providing additional information about the hepatocellular carcinomas originally published in ***Li et al. (2021)***, and Oskar Hallatschek, Chris McFarland and Jona Kayser for useful discussions. This work was supported by an ARCS Fellowship and Big Data for Genomics and Neuroscience Training Grant (MAL), the Howard Hughes Medical Institute (TB), a Swiss National Science Foundation Early Postdoc Mobility Fellowship (NFM), NIH NIGMS R35 GM119774 (TB), the Miller Institute for Basic Research in Science and NIH 1DP2CA280623-01 (AFF). TB is an Investigator of the Howard Hughes Medical Institute.

---

**Figure S1.**
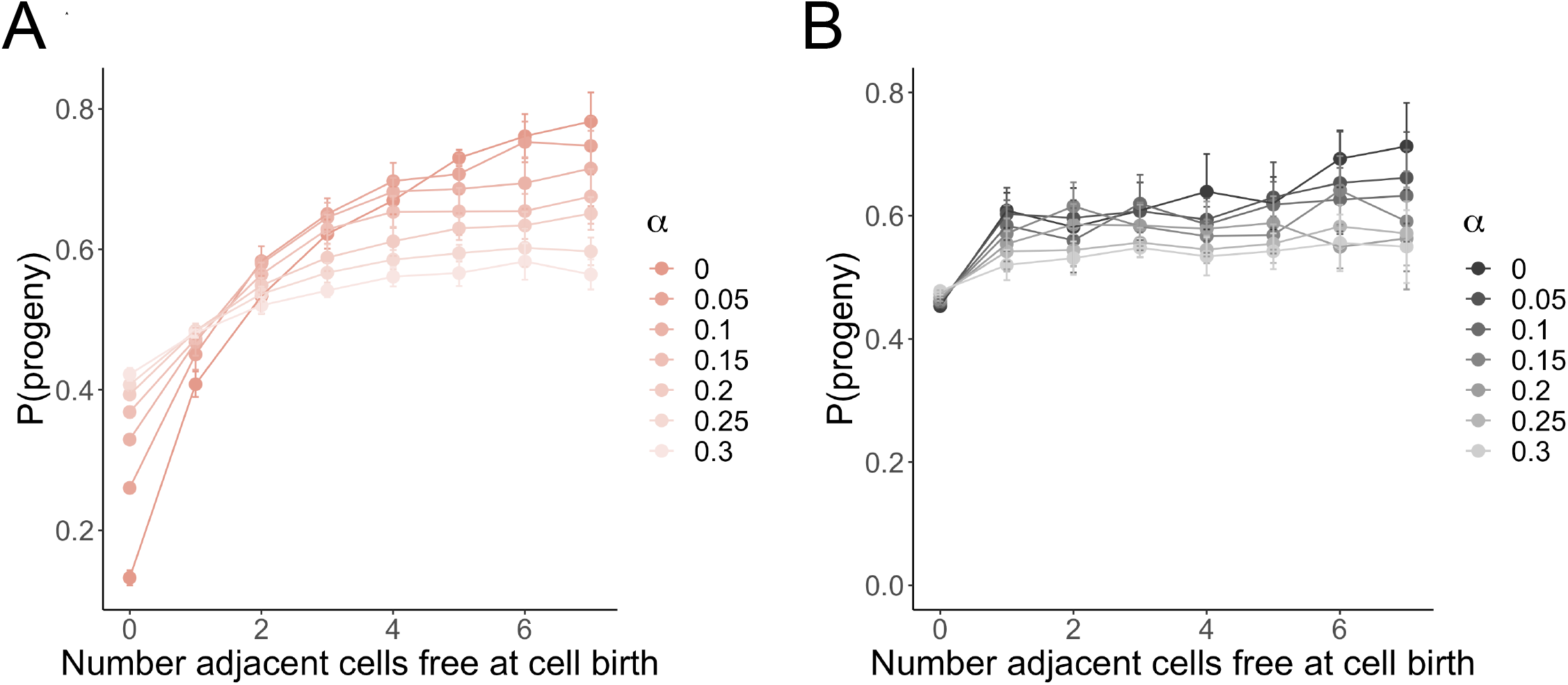
Cellular density creates fitness differences in expanding lattice-based simulations. **A**. Fitness, here approximated by the probability a cell has a daughter cell in the population (P(progeny)) versus the number of adjacent free cells at birth under boundary-driven growth. Spatial impacts on cell fitness are relaxed with increasing cell death rate α (color tint). Means and standard error bars are summarized across 10 simulated tumors per death rate. **B**. Under unrestricted growth, most cells are born into a dense neighborhood (free cells = 0), but fitness is not impacted by spatial location.

**Figure S2.**
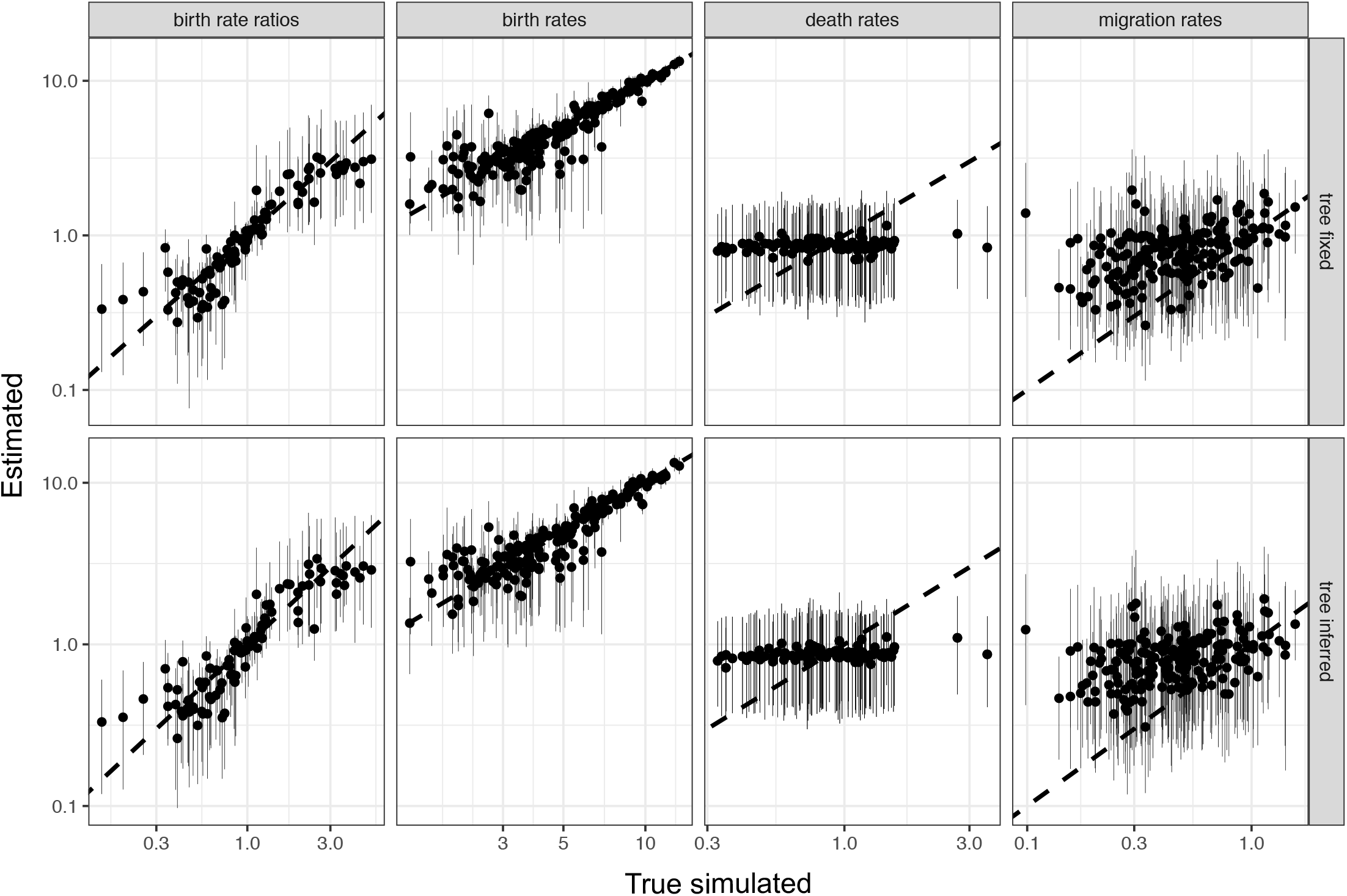
Simulation study to validate SDevo implementation. Birth, death, and transition rates, and ratios of state-dependent birth rates estimated by SDevo versus true population parameters of phylogenetic trees simulated under a two-state birth-death model (see Materials and Methods). Medians (points) and 95% HPD intervals (bars) of estimated values are plotted for each parameter (columns) while either fixing or jointly inferring the tree topology (rows).

**Figure S3.**
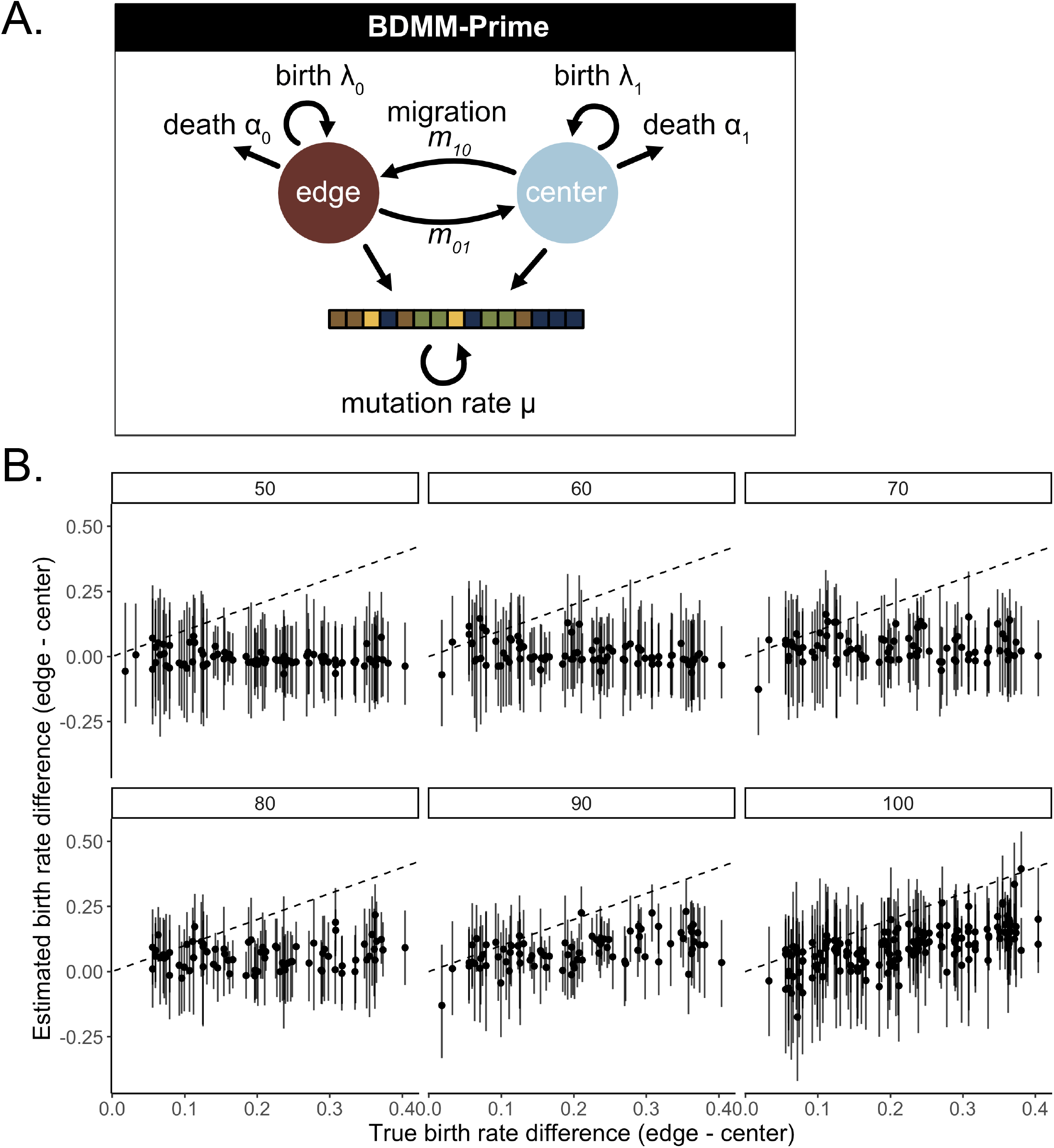
Multi-state diversification models without state-dependent clocks do not sensitively detect growth rate differences in simulated tumors. **A**. Schematic of BDMM-Prime, which does not link state-dependent effects on division to sequence evolution. **B**. True versus estimated means (points) and 95% HPD intervals (bars) of birth rate differences between the edge and center of simulated boundary-driven tumors over a range of sample sizes (headers). Dashed line is *y* = *x*.

**Figure S4.**
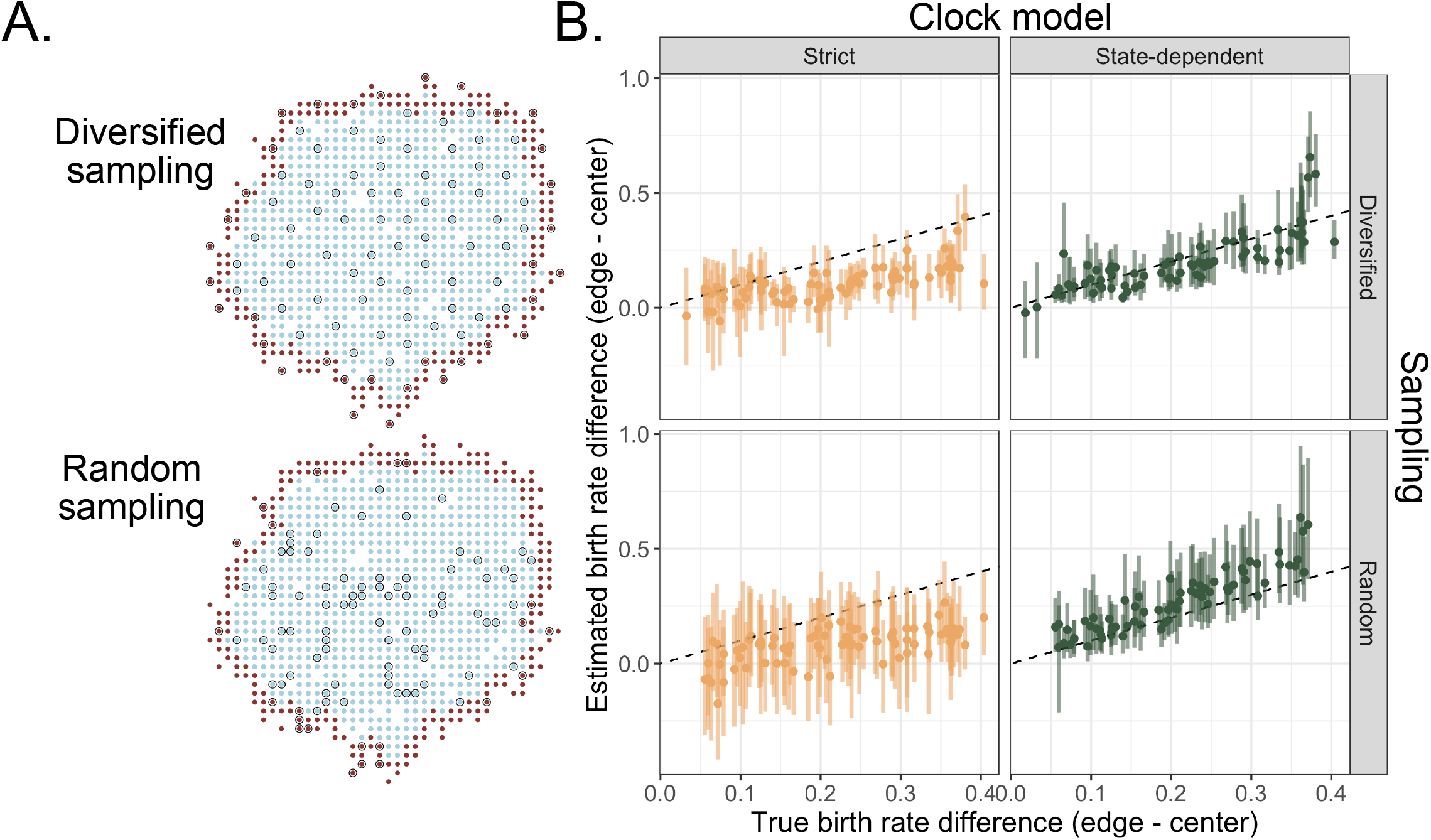
SDevo improves birth rate estimation with more variable (random) sampling over a strict clock model. Example 2D tumors under either diversified or random sampling schemes. Cells are colored by edge (maroon) or center (blue). Grey-highlighted cells are sampled. Diversified sampling maximizes the physical distance between sampled cells. Estimated means (points) and 95% HPD intervals (bars) of birth rate differences between the edge and center of simulated boundary-driven tumors based on 100 sampled cells versus true state-dependent effects (α varies between 0 and 0.435). We compare SDevo (green) with strict clock model (gold) for either diversified or random sampling (rows).2D Growth

**Figure S5.**
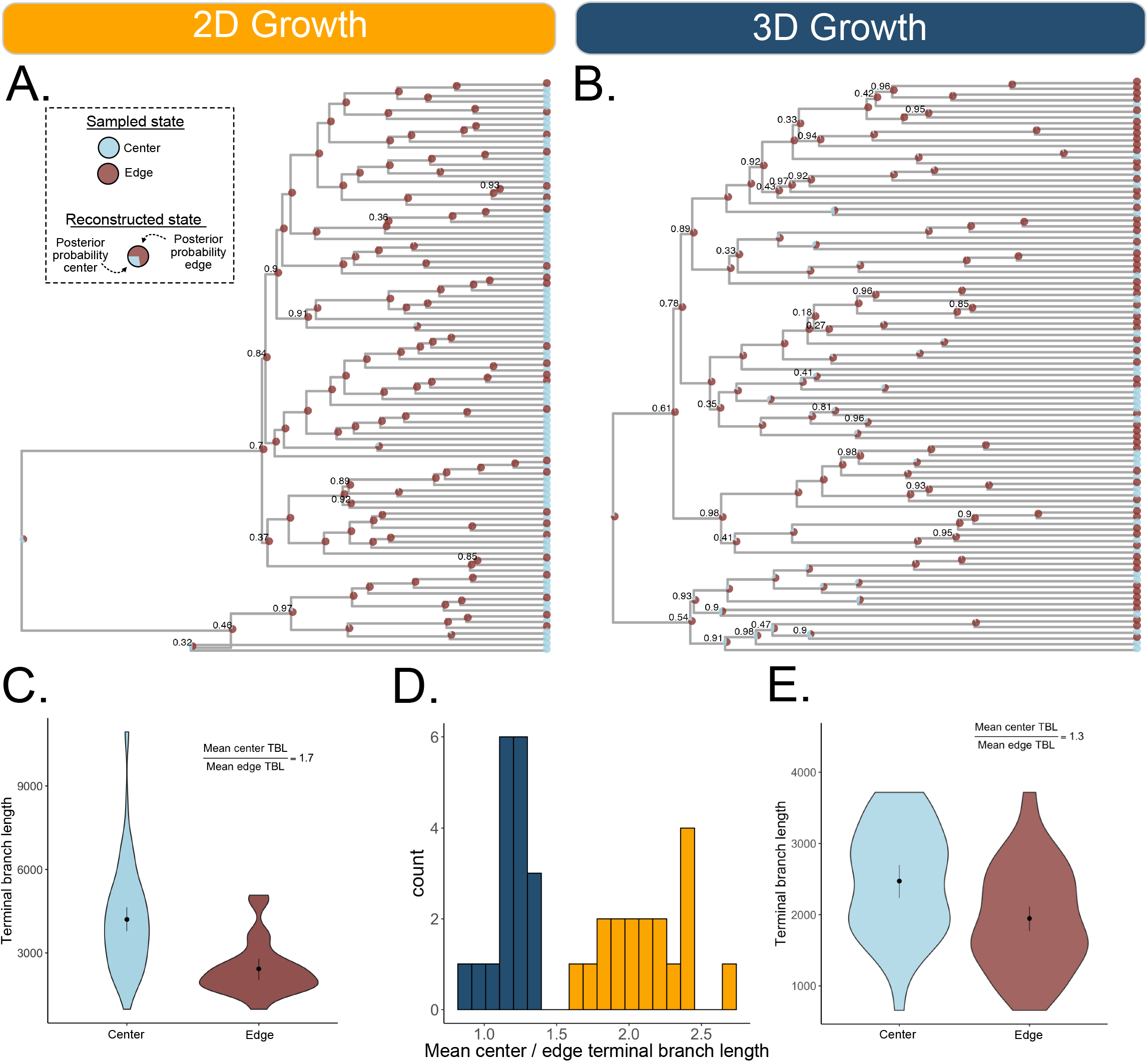
Complex growth and sampling in 3D tumors leads to more variable branching patterns. **A**. Example inferred phylogeny of 2D PhysiCell tumor with reconstructed ancestral edge and center states (*d* = 0.1). Node pie charts represent posterior support for each state. 100 cells were sampled to maximize distance between cells (diversified sampling). **B**. Example inferred phylogeny of 3D PhysiCell tumor with reconstructed ancestral edge and center states (*d* = 0.1). Cells were sampled to maximize distance in 2D space across *z*-slices of the simulated tumor as described in Materials & Methods. For both trees, posterior node support is indicated if less than 99%. **C, E**. Comparisons of inferred terminal branch lengths between cells sampled on the edge and center of 2D and 3D tumors. **D**. Distribution of the relative ratio of center to edge mean terminal branch lengths across multiple simulations with equivalent spatial constraints. Asymmetric branching between edge and center states is observed more often in 2D (gold) than 3D (navy) tumors.

**Figure S6.**
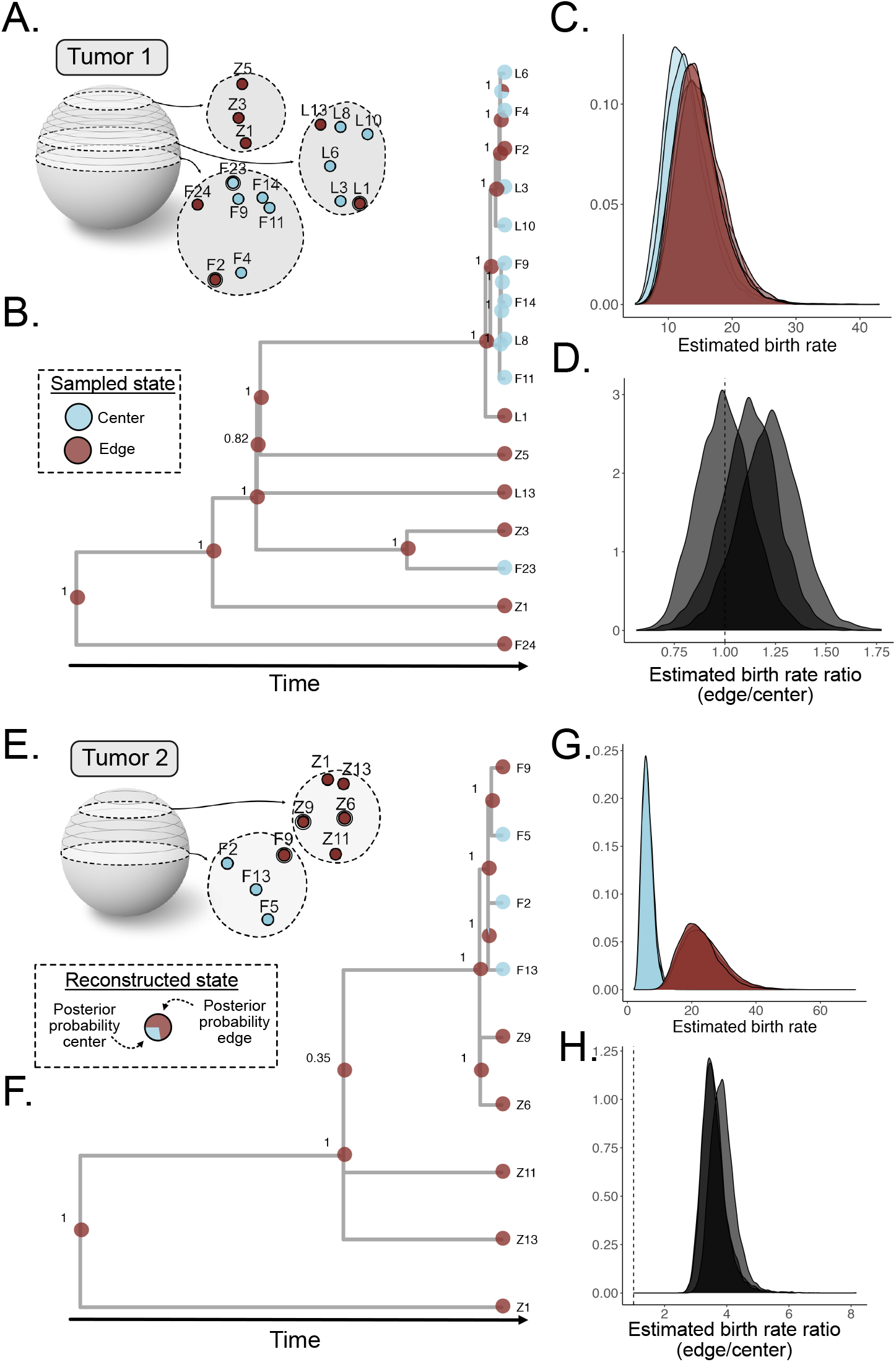
Detection of boundary-driven growth in hepatocellular carcinoma with variation in edge/center state calling. We called an alternate set of states based on a distance of < 10% of each tumor diameter instead of published edge/center labels. **A**. Multi-region sampling map for Tumor 1 adapted from ***Li et al. (2021)*** with alternate state labels. Double outline indicates change in state from published states (Figure 5A). **B**. Inferred tumor phylogeny and reconstructed ancestral spatial states for a single SNV subset. Clade posterior supports are indicated at nodes. **C**. Marginal posterior distributions for edge (maroon) and center (blue) birth rates estimated from the Tumor 1 WGS data across three independent SNV subsets. **D**. Posterior distribution of edge/center birth rate ratio. Dashed line indicates ratio of 1. We estimate a mean 1.11x higher birth rate on the edge compared to center. **E**. Multi-region sampling map with alternate states for Tumor 2 reproduced from ***Li et al. (2021)***. **F**. Tumor 2 tree and ancestral edge/center states inferred from the sampled populations. **G**. Marginal posterior distributions for edge and center estimated birth rates and **H**. edge/center ratio. We37 estimate a mean 3.69x higher birth rate on the edge versus center based on the alternate state calls.

**Figure S7.**
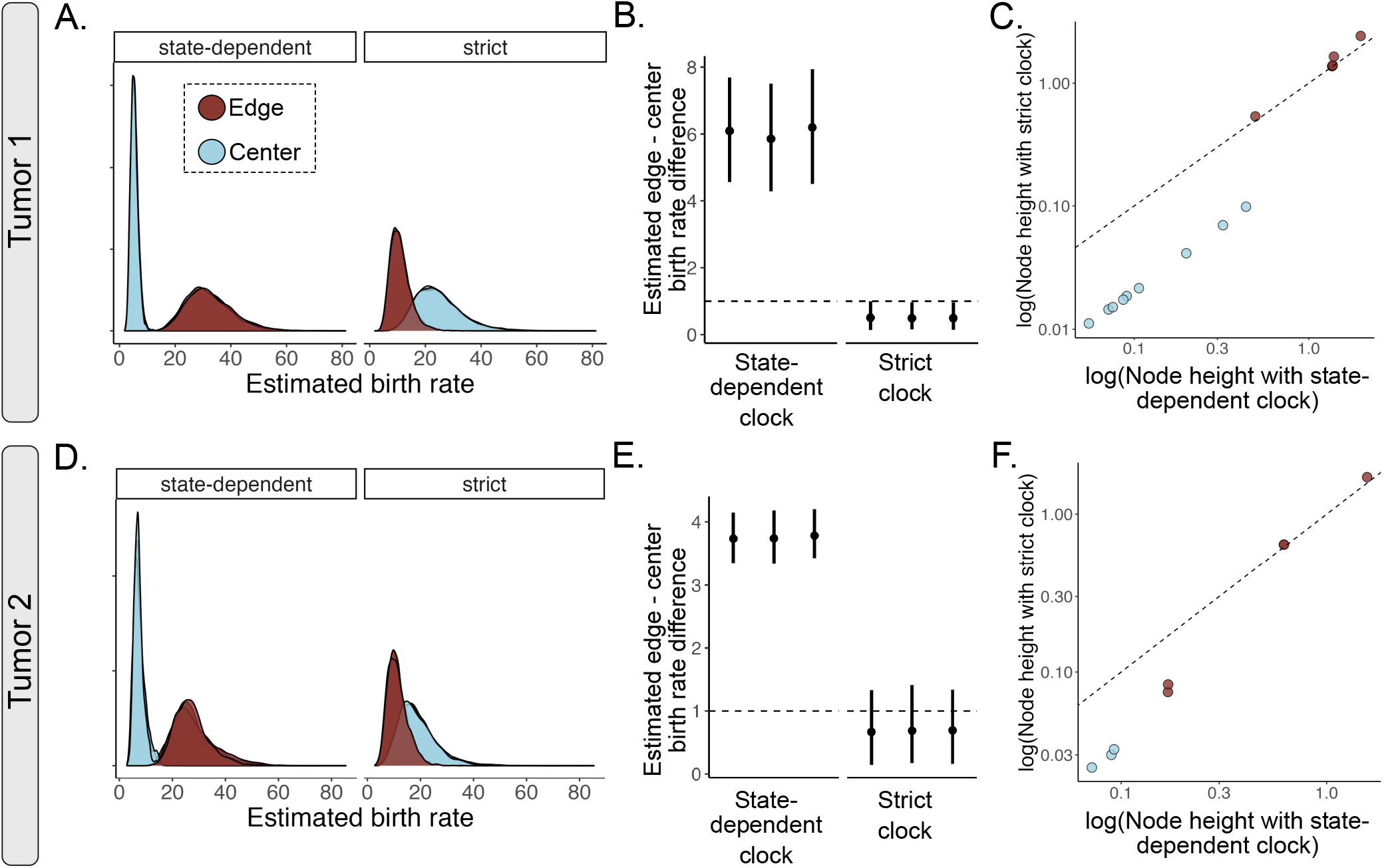
SDevo infers boundary-driven growth in HCC tumors where a strict-clock fails and changes inferred node timings. We compared estimates of birth rate differences between edge and center under a state-dependent birth-death model (BDMM’) using both our novel state-linked sequence evolution model or a strict clock (state-independent) sequence evolution model. For Tumor 1 **(A)**, Tumor 2 **(D)**, posteriors of edge and center birth rate estimates for each sequence evolution model are shown in maroon and blue, respectively. Means and 95% HPD intervals for the inferred birth rate ratios between edge and center states for Tumor 1 **(B)** and Tumor 2 **(E)**. Posteriors are inferred across three independent SNV subsets. Dashed lines indicate ratio of 1. Note, power analyses on simulated tumors (Figure 3 and S3) suggest that the strict clock model should be under-powered and sensitive to sampling variation at these sample sizes (Tumor 1: *n* = 16, Tumor 2: *n* = 9). **C. and F**. Scatterplots show ancestral node heights inferred under strict clock versus heights inferred by SDevo colored by most probable ancestral state. Nodes are compared based on matching a subset of tips.

**Figure S8.**
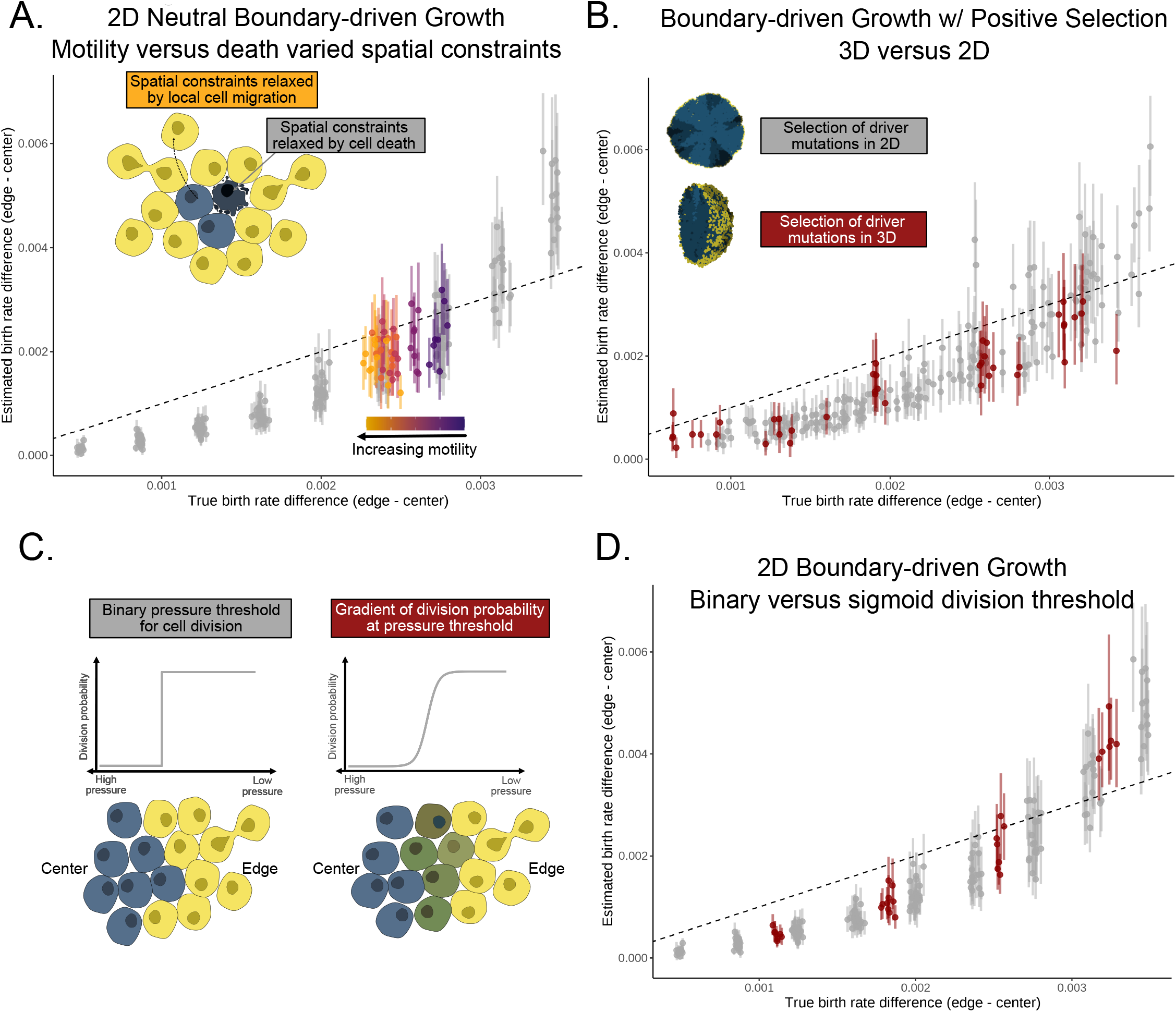
Investigation of extended modes of spatial tumor growth in PhysiCell simulations. **A**. Mean (points) and 95% HPD intervals (bars) of birth rate differences estimated by SDevo when spatial constraints are relaxed by increasing cell motility (purple to orange gradient) compared to when spatial constraints are relaxed by cell death (grey). The x-axis is the effective true birth rate difference in both scenarios. **B**. True versus SDevo-estimated birth rate differences in simulations with both boundary-driven growth and positive selection of driver mutations. We compare simulations in 2D (grey) and 3D (red). **C**. Schematic of simulated relationships between cell pressure and division probability for either a binary (left) or sigmoidal (right) gradient in PhysiCell simulations. D. True versus estimated birth rate differences of simulated tumors with either a binary (grey) or sigmoidal (red) pressure threshold.

**Figure S9.**
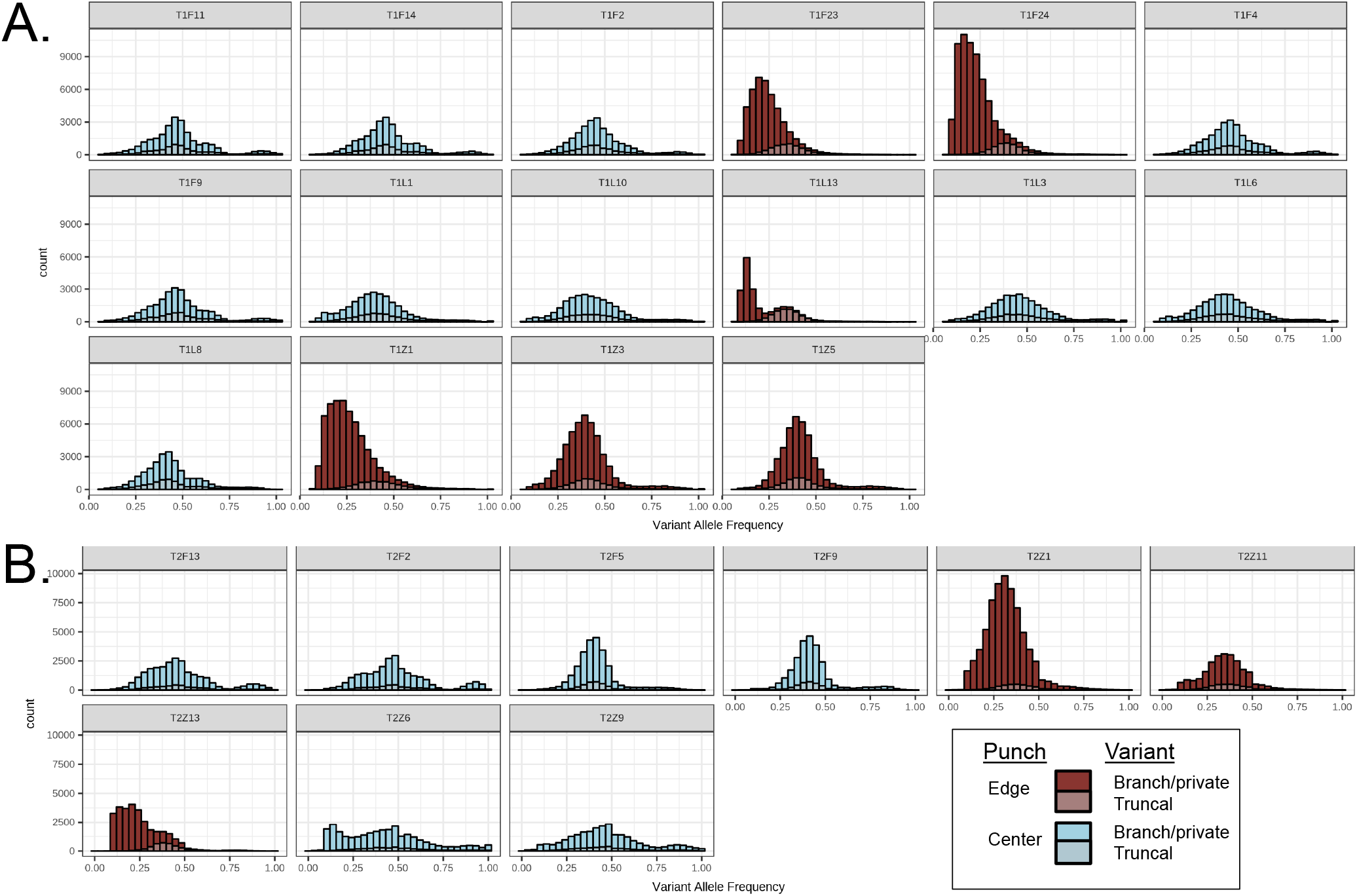
Variant allele frequency (VAF) histograms reveal punches are largely clonal. Variant allele frequencies for all non-truncal (opaque) and truncal (transparent) mutations observed in tumor punches from Tumor 1 (**A**) and Tumor 2 (**B**) reveal that punches contain only a single high frequency clone, with the exception of T1L13. Punches are colored by their edge (maroon) or center (blue) status. State labels correspond to ***Li et al***. (***2021***), Table S8.

**Figure S10.**
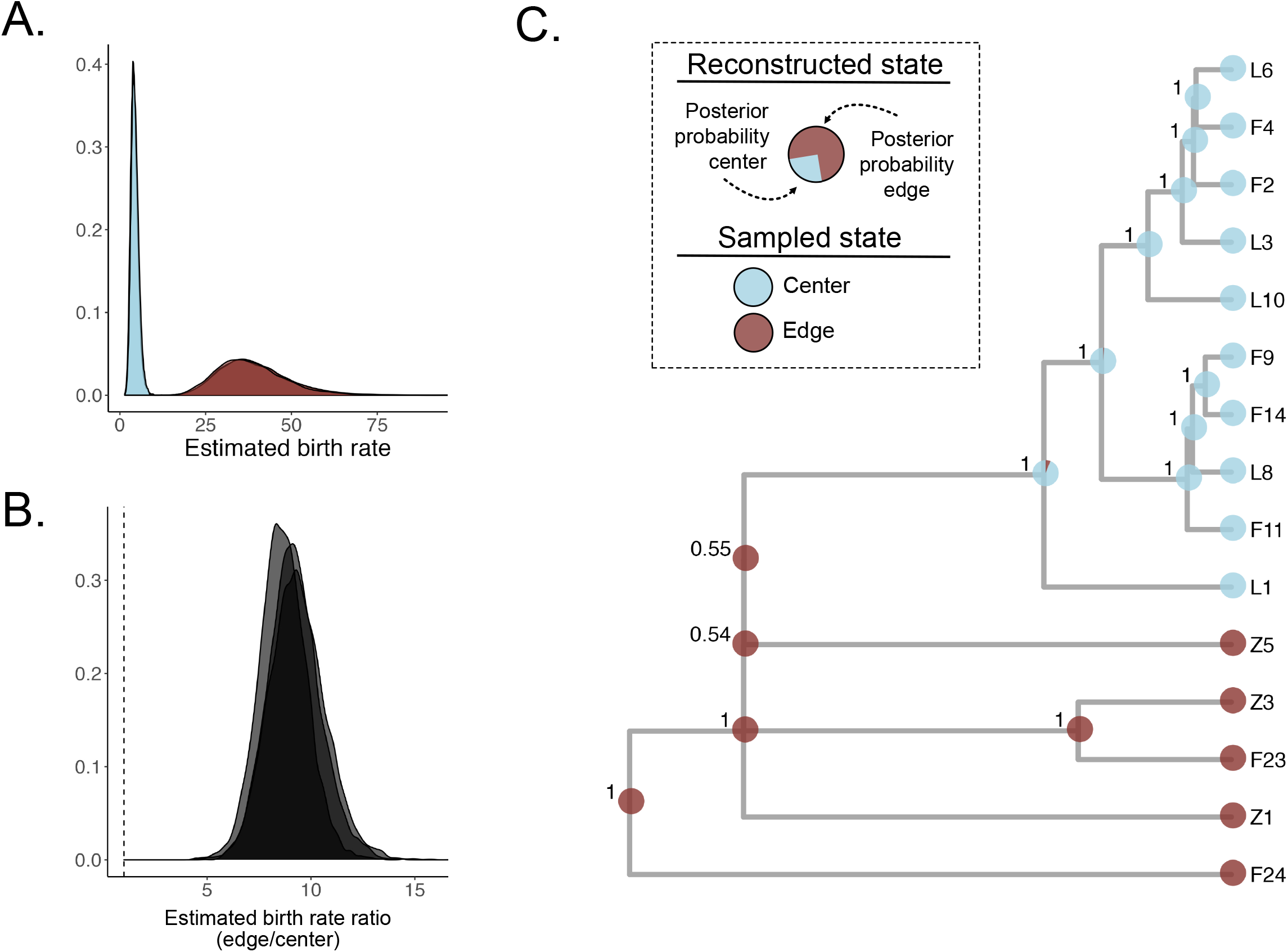
SDevo analysis excluding heterogenous punch T1L13 also estimates birth rate differences between center and edge samples. To ensure our results are not driven by an edge-associated sample (T1L13), which potentially contains multiple subclones, we repeated the analysis of Tumor 1 without this punch. **A**. Marginal posterior distributions for edge (maroon) and center (blue) birth rates estimated from the Tumor 1 WGS data excluding T1L13 inferred for three SNV subsets. **B**. We estimate a 9.06x higher birth rate on the edge compared to center (95% HPD 6.64-11.72x, summarized across the same three independent inferences). **C**. Inferred MCC tumor phylogeny and reconstructed ancestral spatial states. Posterior clade support is indicated at each node.

